# Lack of cortistatin drives neuroimmune and vascular dysfunction in brain ischemia

**DOI:** 10.1101/2024.09.11.611691

**Authors:** Julia Castillo-González, Pablo Vargas-Rodríguez, Ignacio Serrano-Martínez, Alejandro Cuenca-Martagón, Irene Forte-Lago, Melanie Price, Lara Buscemi, Lorenz Hirt, José Luis Ruiz, Elena González-Rey

## Abstract

**Background:** Ischemic stroke remains a leading cause of death and disability, with limited therapies addressing its pathophysiology. Neuroinflammation, blood–brain barrier disruption, and immune dysregulation critically contribute to both acute neuronal injury and chronic functional decline. Elucidating the endogenous factors that govern these processes and understanding the intricate interplay among them is essential for developing effective therapeutic strategies. Cortistatin, a neuropeptide expressed in the nervous and immune systems, exhibits potent immunomodulatory actions, but its role in neuroinflammatory diseases remains unclear.

**Methods:** In this study, we combined human and murine transcriptomic analyses with a preclinical stroke model in cortistatin-deficient mice to investigate the endogenous and therapeutic roles of cortistatin during the acute (48 h) and subacute (7 days) phases of ischemic stroke.

**Results:** We demonstrate for the first time that cortistatin deficiency amplified ischemia-induced transcriptional programs, heightening neuroinflammation and glial/neurovascular dysfunction, and worsening neurological outcomes. Interestingly, cortistatin deficiency accelerated aging-associated pathological features, underscoring its essential role in neuroimmune and cerebrovascular homeostasis. Conversely, cortistatin administration improved neuronal survival, mitigated immune dysfunction, and preserved myelin and vascular integrity during both the acute and subacute stages.

**Conclusions:** These findings provide the first evidence identifying cortistatin as an endogenous neuroimmune regulator in ischemic stroke and a promising multimodal therapeutic agent for improving acute and long-term outcomes.

**Trial registration:** Not applicable

## INTRODUCTION

Ischemic stroke is the second leading cause of mortality worldwide and the main contributor to adult disability [1]. The interruption of cerebral blood flow initiates a cascade of interconnected events, including neuroinflammation, blood-brain barrier (BBB) disruption, and immune deregulation, major hallmarks of stroke pathogenesis [2]. Despite the advancements of recanalization therapies, current treatments remain limited to intravenous recombinant tissue plasminogen activator (rtPA) and endovascular thrombectomy, both with narrow therapeutic window and considerable adverse effects [3]. Therapeutic shortcomings arise from simplistic preclinical models that overlook key risk factors like aging, the use of single-target drugs for a multifactorial disorder, and an overemphasis on immunosuppression rather than immunomodulation. Additionally, the poor understanding of the multi-organ nature of stroke, which involves interactions between the brain, vasculature, and often-overlooked peripheral organs, presents additional challenges [4]. This complex spatiotemporal coordination of immune responses, ranging from acute neuroinflammation to long-term reparative processes, is critical for determining neurological outcome. Consequently, identifying endogenous mediators that modulate neuroimmune and neurovascular homeostasis across both acute and subacute phases of ischemic injury represents a major unmet need and a promising avenue for therapeutic innovation.

Cortistatin is a neuropeptide broadly expressed in the central nervous system (including GABAergic interneurons, microglia, astrocytes, oligodendrocytes, pericytes, and endothelial cells [5–8]) and immune compartments (including macrophages, lymphocytes, monocytes, and dendritic cells [9]), that has emerged as a potent immunomodulatory factor [10,11]. Despite sharing homology, functions, and receptors (SSTR1-5) with somatostatin, cortistatin exerts distinct biological functions also interacting with additional targets, including the ghrelin receptor (GHSR), Mas gene-related receptor X-2, and an unidentified cortistatin-specific receptor [12,13]. Cortistatin has been reported to exert anti-inflammatory and neuroprotective effects across several experimental models of neurological disorders [14–18]. Consistent with these observations, we recently demonstrated a beneficial effect of exogenous cortistatin administration during the acute phase of stroke in young mice [19]. Cortistatin has also been implicated in the regulation of BBB integrity *in vitro* [6], and its deficiency has been associated with increased basal inflammation and heightened vulnerability to inflammatory injury [11,14,16,20].

Despite these findings, the role of endogenous cortistatin in stroke pathophysiology remains unexamined. Specifically, its contribution to neuroimmune crosstalk, BBB stability, and glial responses has not been investigated. Moreover, it is unclear whether cortistatin influences age-related stroke susceptibility or long-term recovery. Addressing these questions represents a novel and unexplored research direction, distinct from prior studies based on exogenous administration or non-ischemic disease models.

Our findings identified cortistatin as a novel endogenous regulator of neuroimmune and neurovascular homeostasis, showing that it restrains maladaptive inflammation while supporting vascular integrity and tissue repair across multiple stages of ischemic injury. Conversely, cortistatin deficiency exacerbates stroke severity and accelerates aging-like neuroimmune and vascular dysfunction, thereby undermining long-term recovery after stroke. Consistent with these findings, human transcriptomic analyses reveal a sustained reduction in cortistatin expression in multiple tissues throughout the course of the disease, underscoring its clinical relevance. Finally, we demonstrated that cortistatin administration reverses key pathological features and improves outcomes across different ages and distinct phases of stroke progression. These findings highlight the potential of targeting endogenous neuroimmune modulators to extend therapeutic opportunities beyond current time-restricted interventions.

## MATERIALS AND METHODS

### Peptides

Lyophilized mouse cortistatin (CST-29) was obtained from Bachem and dissolved in distilled water to a final concentration of 10^-4^ M.

### Animals

The experimental protocols employed in this study were approved by the Animal Care and Use Board and the Ethical Committee of the Spanish National Research Council (Animal Care Unit Committee IPBLN-CSIC # protocol CEEA OCT/2017.EGR) and conducted in accordance with Directive 2010/63/EU of the European Parliament on the protection of animals used for scientific purposes. Transgenic mice deficient in the cortistatin gene (*Cort*^−/−^), generously provided by Dr Luis de Lecea (Stanford University, United States), were bred and maintained on a C57BL/6J background as previously described [12]. Mice were housed in groups of ten per cage under controlled environmental conditions (22 ± 1°C, 60-70% relative humidity) with a 12h light/dark cycle (lights on at 7:00 a.m.), with *ad libitum* access to food and water.

### Transient middle cerebral artery occlusion model

3-month-old and 6-month-old *Cort*^+/+^ and *Cort*^−/−^ male mice were evaluated. Animals from both ages were subjected to the MCAO model as previously described [19]. This model is a robust and highly reproducible gold-standard technique, rigorously validated in stroke research for over two decades [21–24]. It remains the most clinically relevant surgical approach to mimic human pathophysiology, also offering superior controllability of reperfusion and no craniotomy [21]. In MCAO, blood flow is interrupted proximal to the lenticulostriate arteries, making the striatum (caudate-putamen) exceptionally vulnerable to ischemic insult. Consequently, the infarct consistently originates in this core territory and, depending on the severity of the stroke or the susceptibility of the cohorts, the damage can expand from the striatum into cortical areas. For MCAO, anaesthesia was induced using isoflurane (1.5-2% in 70% N_2_O/30% O_2_) via a facemask, and meloxicam (2mg/kg, Norbrook) was administered subcutaneously for post-surgery analgesia. Body temperature was maintained constant throughout the procedure using a heating pad. Regional cerebral blood flow (rCBF) was continuously monitored by laser Doppler flowmetry (moorVMS-LDF1, Moor Instruments) fixing a probe on the skull over the core area supplied by the middle cerebral artery (MCA) (approximately 1 mm posterior and 6 mm lateral to the bregma). Transient focal ischemia was induced by introducing a silicon-coated filament (701756PK5Re, Doccol) through the left common carotid artery into the internal carotid artery to occlude the MCA. The surgery was considered successful if the rCBF decreased below 20% of the baseline during occlusion and recovered above 50% of the baseline upon reperfusion. Following surgery, mice were returned to their cages with *ad libitum* access to food and water. Each cage was supplied with water-softened chow and recovery gel. To ensure an equivalent ischemic insult across age groups despite age-related vascular differences, the occlusion was maintained for 20 min in 3-month-old mice and 35 min in 6-month-old mice, using a standardized filament size for both cohorts. After this time, the filament was withdrawn to allow reperfusion. While longer occlusion times are frequently reported, we implemented a refined, non-lethal stroke model to minimize mortality (*i.e.,* thinner intraluminal filaments yields smaller and more reproducible infarct sizes, reduces endothelial damage, increases reperfusion rates [19,21]) and allow a reliable assessment of long-term recovery mechanisms. Moreover, shorter occlusion durations have been shown to produce infarct volumes comparable to over half of stroke patients [25–27].

Cortistatin was administered intraperitoneally at a standardized dose of 144 µg/kg, a regimen selected based on our previous optimization studies identifying the optimal therapeutic window for neuroprotection [19], and consistent with its established efficacy in other neuroinflammatory and autoimmune models [16,20]. Treatment was initiated 24 h post-ischemia and delivered either as a single dose (acute phase) or as daily injections for 7 consecutive days (subacute phase). Control groups received equivalent volumes of saline solution. Neurological assessments, behavioural tests, ischemic volume determination, morphometric characterization of glial population, and inflammatory response evaluation were determined at 48 h and 7 days after surgery. We focused on 48 h post-stroke to study the potential endogenous and therapeutic roles of cortistatin because it represents a critical period for key processes, including neuronal damage, glial activation, BBB disruption, and immune modulation [1,28,29]. The 7-day post-ischemia interval was selected to capture early long-term recovery phase, a period characterized by remodelling processes supporting active plasticity and repair mechanisms and therapeutic responsiveness [30,31].

### Neurological and behavioural assessments

The functional outcome of each mouse was evaluated using a neurological score every 24 h following MCAO and prior to sacrifice. Neurological severity was graded on a scale from 0 to 3, assessing motor, sensory, reflex, and balance functions, based on a combination of established scoring systems [32].

In addition, muscle coordination and endurance were analysed using the wire-hanging test. Mice were suspended on a custom-made apparatus consisting of a single wire stretched between two posts over a soft surface. Training was conducted 24 h before surgery, and performance was evaluated at 48 h and 5 days post-MCAO, recording the number of successful reaches (defined as the mouse touching either the right or left post) or falls [33]. The test concluded after 180 s or when a mouse fell from the wire ten times. Motor skills were further characterized using the pole test, which measured the ability of mice to grip and descend a vertical pole to a safe surface [34]. Mice underwent one training trial 24 h before surgery. Post-surgical evaluations of mice were conducted 48 h and 5 days after MCAO. Performance was scored based on successful descents or falls (reaching the mat or falling onto it, respectively). The total time taken to reach the mat with at least two limbs, along with the number of turns required to descend, were recorded. The test concluded after three trials, if any trial exceeded 120 s, or after three falls. Cognitive function was evaluated using the novel object recognition (NOR) test to assess long-term episodic-like memory. Behavioural performance was recorded for 10 min using a camera and analysed with automated software (SMART V3.0.06, Panlab). Prior to testing, mice underwent a 15-min habituation period in the arena. Baseline cognitive capacity was established starting 24 h before the stroke procedure; mice were first exposed to two identical objects, followed 24 h later (30 min before the stroke) by a session where one object was replaced with a novel one. Following the stroke, the 7-day memory assessment was conducted: 6 days post-stroke, mice were again exposed to two identical objects, and 24 hours later (day 7), one object was replaced by a novel object. Exploration was defined as the animal’s snout being directed toward the object within a distance of 2 cm. To quantify memory performance, the recognition index (RI) and discrimination index (DI) were calculated. The RI was defined as the time spent exploring the novel object divided by the total exploration time (T-novel / [T-novel + T-familiar]). The DI was calculated as the difference in exploration time between the novel and familiar objects divided by the total exploration time ([T-novel – T-familiar] / [T-novel + T-familiar]). Values approaching 1 reflect a stronger and more robust memory performance.

### Ischemic lesion volume determination

At 48 h or 7 days post-MCAO, mice were sacrificed by an overdose of intraperitoneal ketamine (100 mg/kg, Ritcher Pharma) and xylazine (10 mg/kg, Fatro Iberica), followed by intracardial perfusion with 4% PFA. Brains were dissected, post-fixed in 4% PFA for 24 h at 4°C, and then cryoprotected in 30% sucrose for 48 h at 4°C. Subsequently, brains were frozen in isopentane on dry ice at –25°C and stored at –80°C until further processing. Coronal brain sections were cut at a thickness of 20 µm using a cryostat.

To assess tissue damage and quantify infarct volume, serial sections were stained with cresyl violet (CV), which is the most appropriate tool at 48h post-stroke and beyond. This basic dye binds to neuronal RNA, causing healthy tissue to stain dark purple while the infarcted core appears pale or white due to the loss of cellular RNA and neuronal lysis. The procedure followed a protocol of gradual hydration in ethanol, staining with 1 mg/ml of CV-acetate in distilled water, gradual dehydration in ethanol, and clearing in xylene [32]. Stained sections were imaged using a Leica DM2000 Transmit stereomicroscope at 2.5x magnification, and infarcted areas were measure using the ImageJ Fiji free software. Infarct volume was calculated by summing the infarcted areas from each section and multiplying by the spacing distance between sections.

### Immunofluorescence

Free-floating brain sections (20 µm-thick) were first washed with PBS, then blocked and permeabilized using a solution of PBS 1x, 1% bovine serum albumin (BSA), 0.1% Triton X-100, and 5% goat serum for 60 min at RT. The sections were subsequently incubated overnight at 4°C with primary antibodies anti-MAP2 (1:1,000, Sigma MAB3418), anti-Iba1 (1:1,000, Wako 19-19741), anti-GFAP (1:5,000, Dako 20334), anti-CD31/PECAM-1 (1:300, BD550274), anti-ZO-1 (1:100, Fisher 40-2200), anti-VEGF (1:500, Abcam 52917) or APC-labelled anti-CD45 (1:200, BD 559864) diluted in PBS 1x with 1% BSA, 0.1% Triton X-100, and 5% goat-serum. Next, sections were washed with PBS and then incubated with the appropriate Alexa Fluor-conjugated secondary antibodies (1:1,000, Sigma) and DAPI in PBS 1x, 1% BSA, 0.1% Triton X-100, and 2% goat-serum for 60 min. Sections were mounted using Mowiol. For IgG staining, the same protocol was followed, except that an Alexa Fluor 594-conjugated goat anti-mouse IgG antibody was used without a primary antibody. Images were obtained with a LEICA DMi8 S Platform Life-cell microscope at 20x magnification and a Leica TCS SP8 STED confocal microscope at 63x magnification.

### Morphometric analyses of microglial phenotypes

To thoroughly characterize the complex and heterogeneous morphology of microglia/infiltrating macrophages following stroke, we performed extensive morphometric analyses. First, individual microglia/macrophages from specific brain regions, such as the periinfarct area, were randomly selected. Attention was warranted to avoid cell overlap and to ensure complete capture of both the soma nucleus and cellular branches. A total of 15-30 cells per mouse from 3-6 mice (N=3-6) were analysed. To ensure objectivity and eliminate observer bias, image processing was performed using ImageJ Fiji software, following a systematic, blinded, and automated protocol adapted from previous studies [17,35,36].

Initially, images were converted to 8-bit grayscale and automatically pre-processed to reduce background noise and enhance contrast. Subsequently, images were binarized using a pre-determined threshold applied consistently across all experimental groups, and non-cellular artefacts were removed using the *remove outliers* and *despeckle* functions. Manual refinement was strictly limited to the removal of non-cellular artifacts and debris to ensure the accuracy of the segmentation. After image processing, morphometric parameters were obtained automatically using the Skeletonize and Fractal Analysis plugins in ImageJ Fiji [37]. For *skeleton analysis*, the number of junctions and maximum branch length were quantified. For *fractal analysis*, key parameters included the convex hull area (the minimal convex polygon enclosing the entire cell structure), span ratio (defined as the ratio of the major to minor axes of the convex hull), and fractal dimension (an index representing cellular complexity).

### Isolation of adult microglia and astrocytes

Adult brains from 3-month-old *Cort+/+* and *Cort−/−* male mice were harvested under sterile conditions and dissociated into single-cell suspensions using the Adult Brain Dissociation Kit (Miltenyi Biotec) according to the manufacturer’s instructions [38]. Briefly, after enzymatic and mechanical dissociation with the gentleMACS Dissociator, myelin, debris, and erythrocytes were removed to obtain a clean cell suspension that was used as a mixed neuron-glial culture or for obtaining individual microglia and astrocytes. Microglia were isolated by magnetic separation following enrichment with CD11b MicroBeads, whereas astrocytes were isolated using the Anti-ACSA-2 MicroBead kit. When required, Fc receptor blocking reagent was included to reduce nonspecific binding. Purity and viability of the isolated populations were verified by immunocytochemistry using cell type-specific markers. The primary mixed neuron-glial cocultures consisted of 17 ± 5% neurons, 44 ± 2.5% astrocytes, 30 ± 2.5% oligodendrocytes, and 9 ± 1.5% microglia (mean ± SEM).

### LPS and oxygen-glucose deprivation and reoxygenation exposure

Isolated mixed neuron-glia, microglia and astrocytes were plated at 10⁵ cells/well in 24-well plates coated with poly-D-lysine (5 µg/mL) under standard culture conditions and allowed to recover before experimental manipulation. To simulate an inflammatory environment, LPS from *E. coli* 055:B5 (100 ng/ml, Sigma) was added to the cells for 24 h, after which supernatants were collected. To model oxygen-glucose deprivation and reoxygenation (OGD/R), cells were incubated in glucose-free medium under hypoxic conditions for the deprivation phase (2 h for microglia/astrocytes; 4 h for neuron-glial cultures) and subsequently returned to normoxic, glucose-containing culture conditions for the reoxygenation period (20-22 h), in the presence/absence of exogenous cortistatin (100 nM). To model ischemia-associated inflammatory activation, after OGD/R for the indicated period (2/4 h + 22/20 h), cells were washed and then incubated in fresh medium supplemented with LPS (100 ng/ml, Sigma) in the presence or absence of exogenous cortistatin (100 nM) for another 24 h. Following LPS incubation, supernatants were collected, and inflammatory factors were evaluated by ELISA (as described below). Nitric oxide levels were quantified by the Griess reaction to evaluate the cellular redox state as a complementary endpoint.

### Determination of the peripheral immune response

To characterize the peripheral immune activity, we evaluated the levels of immune mediators and the spleen response. To quantify peripheral inflammatory factors, blood samples were collected by cardiac puncture, allowed to clot for 2 h at RT, and then centrifuged at 2,000 rpm for 10 min. The concentrations of the factors in serum or in supernatants from the *in vitro* cultures above were measured using a sandwich ELISA assay. Maxisorb plates (Millipore) were coated overnight at 4°C with capture antibodies (anti-IL-6, anti-IL-12, anti-TNF-α from BD Pharmigen, or anti-MCP-1 from PeproTech). Plates were washed with PBS containing 0.05% Tween-20 and then blocked with PBS containing 10% fetal bovine serum (FBS) for 2 h. Serum samples and recombinant standards were added to the plates and incubated overnight at 4°C. After incubation, biotinylated detection antibodies (Serotec) were applied for 1 h at RT. Following washing, plates were incubated with avidin-HRP (Sigma) (1:500 in blocking buffer) for 1 h and developed using ABTS substrate (Sigma). Absorbance was measured at 405 nm with a spectrophotometer (Molecular Devices), and the concentrations of inflammatory factors were calculated based on the standard curve generated concurrently.

Spleens were collected, weighed, and gently dissociated into a 100 mm culture dish. The homogenized tissue was transferred into RPMI medium supplemented with 10% FBS, 2 mM L-glutamine, 1% P/S, and 50 µM 2-mercaptoethanol. The mixture was then let to sit on ice for 5 min, and the cell suspension was carefully collected and centrifuged at 1,670 rpm for 10 min at 4°C. The resulting pellet was resuspended in Ack lysis buffer (150 mM NH_4_Cl, 10 mM KHCO_3_, 0.1 mM Na_2_EDTA) for 10 min at RT to lyse red blood cells. RPMI medium was added, and the cells were centrifuged again at 1,670 rpm for 10 min at RT before being counted.

### RNA extraction and evaluation of gene expression

After MCAO, brains were collected at the specified time points (*i.e.,* 48 h and 7 days) and separated into contralateral and ipsilateral hemispheres. For RNA isolation, brain tissue was placed in TriPure reagent (Roche). cDNA synthesis from 20 ng RNA per sample was performed using the RevertAid First Strand cDNA Synthesis Kit (ThermoFisher) with random hexamers in a CFX Connect QPCR System (Biorad) under the following conditions: incubation at 25°C for 5 min, reverse transcription at 42°C for 60 min, inactivation at 70°C for 5 min. Gene expression analysis was performed using conventional qPCR. cDNA was added to the SensiFast Sybr No-Rox kit 1x (Bioscience), along with forward and reverse primers specific to each gene and nuclease-free water to reach a final volume of 20 µl. The qPCR reactions were run in a CFX96 thermocycler (BioRad) under the following conditions: initial polymerase activation at 94°C for 5 min, 40 cycles at 94°C for 30 s (denaturation), 58°C/60°C/62°C for 30 s (primer-specific annealing), and 72°C for 30 s (extension). Gene expression quantification was performed using the comparative threshold cycle (Ct) method, with Ct values normalized to those of the housekeeping gene *Rplp0*. Fold change was calculated using the ΔΔCt method. The specific primers for each gene are listed in Additional File 1: Table S1.

### Next-generation transcriptome sequencing (RNA-seq) and data analysis

RNA (200 ng) from contralateral (CL, healthy) and ischemic (MCAO) brain hemispheres isolated from *Cort^+/+^* and *Cort^−/−^*mice 48 h post-stroke was used to prepare mRNA libraries with the Illumina stranded mRNA Prep Ligation kit (Illumina, San Diego, CA, USA). Three (CL) and five (MCAO) independent biological replicates were used with RNA Integrity Number coefficients >8.7 (Bioanalyzer RNA 6000 Nano-chip, Agilent). Quality and size distribution of mRNA libraries were validated using the Bioanalyzer High Sensitivity DNA assay, and their concentrations were measured on a Qubit fluorometer (ThermoFisher). Final libraries were pooled in equimolecular ratios and converted to single-stranded circular DNA with MGIEasy Universal Library Conversion kit (MGI Tech, Shenzhen) for sequencing on MGI platforms. The 100 bp paired-end sequencing was conducted on a DNBSEQ-G400 sequencer (MGI Tech, Shenzhen). The average percentage of sequencing reads above a quality score of Q>30 was 96.3%, with a mean GC content of 50.0%, and over 44 million reads per sample.

We implemented the reanalyzerGSE v3.0.0 software [39] in the RNA-seq analysis to identify differentially expressed genes (DEGs) and to prevent the exclusion of lowly expressed genes with potential biological relevance, such as cortistatin (see details in Additional File 2). We detected >45,000 transcripts. Multidimensional scaling (Principal Coordinates Analysis, PCoA) and unsupervised hierarchical clustering of normalized samples were performed to assess the divergence and the replicability of samples from each independent comparison. DEGs with a False Discovery Rate (FDR) <0.05 were calculated by comparing: i) *Cort^−/−^ vs. Cort^+/+^* samples for each condition (CL and MCAO); ii) MCAO *vs.* CL for each genotype (*Cort^−/−^* and *Cort^+/+^*). The log2 fold change (log_2_FC) was used to quantify gene expression changes. To explore the functional significance of DEGs, we performed two different approaches: an over-representation analysis based on the Gene Ontology (GO) database [40,41] and a Gene Set Enrichment Analysis (GSEA), as indicated in Additional File 2. Conventional qPCR was used to validate differentially expressed genes selected from our transcriptomic studies.

### Cell-specific gene sets and cell type proportion estimation

Cell-type–specific gene sets were obtained from both the default settings of the markerGeneProfile R package v1.0.4 [42] and the curated marker sets reported by Zheng et al [43]. Both sources include overlapping gene markers for microglia, astrocytes, and oligodendrocytes; however, the astrocyte and oligodendrocyte gene sets differ in composition and biological meaning between the two approaches. Unlike other marker databases that rely on a single data source, these marker sets represent a consensus from several published studies and account for brain regional heterogeneity. We use both gene sets to estimate marker gene profiles (MGPs), which serve as a proxy for relative changes in cell type proportions. Importantly, because marker gene expression can fluctuate significantly across different cellular states, this approach was not intended to quantify absolute cell populations, but instead to be used strictly as an estimation of the relative transcriptomic footprint and marker enrichment of these cell populations within the bulk tissue. For this, we used as input the transcript-per-million (TPM) normalized gene expression data from our RNA-seq. Details on how MGPs were derived can be found in Additional File 2. Both gene sets were also used to assess the enrichment of our samples for the specified cell markers by performing a GSEA as a complementary analysis.

### Integration of human and mouse datasets for cortistatin gene expression analysis

Cortistatin gene expression levels were obtained from the Gene Expression Omnibus database (GEO), incorporating previously published datasets from human and mouse samples following stroke. For humans, we reanalysed data from: peripheral blood mononuclear cells (PBMCs) from ischemic patients at acute and subacute stroke stages (GSE122709) [44]; and with/without haemorrhagic transformation (GSE199435) [45]; leukocytes from patients with differential stroke severity (GSE202709) [46]; and brain tissue dissected from healthy controls and acute ischemic stroke patients post-mortem (GSE56267) [47]. Mouse data from cortical tissue were reanalysed from different experimental stroke models, such as the MCAO model (GSE253799, unpublished; GSE227186 [48]); cerebral venous sinus thrombosis (CVST) model (GSE245455, unpublished); and photothrombotic (PT) stroke model (GSE261613, GSE184917 [49,50]; GSE280846, unpublished). Details about patient characteristics, stroke subtype, or time post-stroke can be found in Additional File 2.

### Ethical approval and consent to participate

This study involves the re-analysis of publicly available human data. The original studies from which the data were obtained received ethical approval from their respective Institutional Review Boards (see references above). Therefore, a new ethical approval or informed consent was not required for this manuscript, in accordance with local legislation and institutional requirements regarding the secondary use of anonymous public data.

### Clinical trial number

Not applicable

### Statistical analysis

Data are expressed as the mean ± SEM. Randomization of animals was performed using specialized software (*e.g.,* Research Randomizer). All experiments were conducted in a blinded manner. The number of independent animals (N), as well as the number of sections, cells, or structures analysed (n), are indicated for each experiment. Comparisons between two groups were analysed using the unpaired Student’s t test for parametric data with normal distribution or the Mann-Whitney U-test for non-parametric data. For comparisons involving three or more groups with normally distributed parametric data, one-way ANOVA with Fisheŕs LSD *post-hoc* test was employed. For non-parametric data involving three or more groups, the Kruskal-Wallis test with a Dunn *post-hoc* test was used. When standard deviations were assumed to be different, Brown-Forsythe and Welch ANOVA tests were applied, followed by a Dunnett T3 *post-hoc* test. Spearman’s rho non-parametric test was used for correlation studies. All analyses were performed using GraphPad Prism v8.3.0 software. For the statistical approaches used in the sequencing data analyses, see the section above. We considered *p*-values <0.05 as significant.

## RESULTS

### Cortistatin is downregulated in ischemic stroke

To investigate the potential involvement of cortistatin in ischemic stroke, we first assessed its expression levels in human samples and murine models reanalysing previously published cerebral ischemia datasets. Due to the limited availability of post-mortem human brain tissue and murine blood samples, we primarily used datasets of PBMCs and leukocytes from patients and mouse brain tissue. Our results show that cortistatin expression was markedly reduced in PBMCs in acute and subacute stroke phases and was inversely associated with stroke severity (Fig. 1A). Reanalysis of post-mortem human brain samples revealed a downward trend in cortistatin levels after ischemia (Fig. 1A). Similarly, mouse brain tissue exhibited significant cortistatin downregulation across multiple stroke paradigms, including transient MCAO, CVST, and PT (Fig.1B). Notably, data from the transient MCAO model used in this study, which recapitulates key features of human ischemic condition [51], also resulted in cortistatin downregulation, which was partially reversed by exogenous addition of the neuropeptide (Fig. 1C). These findings implicate cortistatin downregulation as a consistent feature of ischemic stroke pathology.

**Figure 1.**
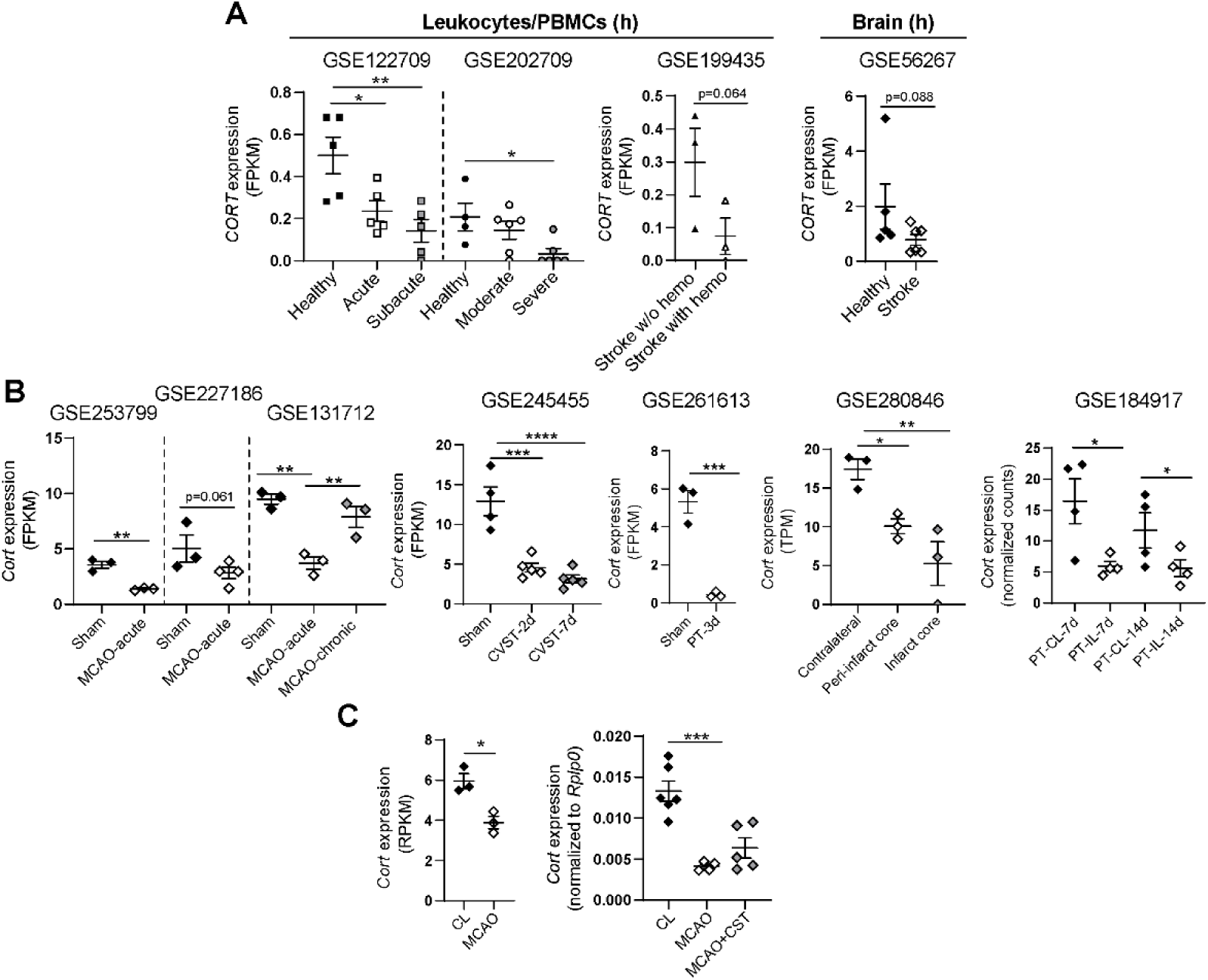
Cortistatin gene expression is downregulated following ischemic stroke in both human and mouse. Cortistatin gene expression levels (normalized RPKM, FPKM, TPM, or count values) obtained from the Gene Expression Omnibus (GEO) database. **(A)** Left, human PBMCs from healthy individuals and ischemic stroke patients at acute (24 h, N=5) and subacute (7 days, N=5) stages (GSE122709); peripheral leukocytes from patients with moderate (NIHSS score<8, N=6) and severe (NIHSS score≥8, N=6) ischemic stroke, compared to healthy controls (N=4) (GSE202709); and PBMCs from ischemic stroke patients with large artery occlusion undergoing thrombectomy, with/without subsequent intracranial haemorrhage (N=3) (GSE199435). Right, cortical tissue of deceased ischemic stroke patients, using matched contralateral (CL) tissue as control (N=5) (GSE56267). **(B)** Cortex from *sham* and ischemic mice (60 min MCAO, sacrificed at 24 h) (GSE253799: peri-infarct cortex and GSE227186: peri-lesion, N=3 each), and ipsilateral (IL) cortex from *sham*/MCAO mice (60 min) sacrificed at 24 h (acute phase) or 28 days post-stroke (chronic phase) (GSE131712, N=3). Cortex of sham (N=4) and mice that underwent cerebral venous sinus thrombosis (CVST), 2– and 7-days post-injury (N=5 each) (GSE245455). Photothrombotic (PT) models: *sham* and peri-infarct cortex 3 days post-injury (GSE261613, N=3); contralateral, peri-infarct and infarct core regions 7 days post-injury (GSE280846, N=3); contralateral and peri-infarct cortex at 7– and 14-days post-stroke (GSE184917, N=4). **(C)** Left, contralateral/ipsilateral hemispheres of MCAO mice (20 min, sacrificed at 48h), analysed by RNA-seq in this study. Right, qPCR validation normalized to *Rplp0*. Data are expressed as mean ± SEM, dots represent individual mice/patients. Statistical analyses were performed using unpaired t-tests or one-way ANOVA for comparisons among multiple groups. \**p≤*0.05, \*\**p≤*0.01, \*\*\**p*≤0.001, \*\*\*\**p*≤0.0001. CST: cortistatin.

### Deficiency in cortistatin enhances immune activity, vascular disorganization, and neurodegenerative responses following ischemic stroke

Given that lower cortistatin gene expression was associated with more severe strokes, we investigated whether cortistatin deficiency plays a role in stroke pathophysiology by analysing the transcriptomic profiles of both cortistatin-deficient (*Cort*^−/−^) and wild-type mice (*Cort*^+/+^) subjected to mild MCAO.

Comparative analysis revealed minimal baseline transcriptomic differences in non-ischemic hemispheres, with 25 DEGs (15 upregulated, 10 downregulated in *Cort*^−/−^ mice; FDR<0.05) (Table S2; Fig. S1A). Similarly, ischemic hemispheres displayed 27 DEGs between genotypes (12 upregulated, 15 downregulated in *Cort*^−/−^ mice), most of which were common across ischemic and healthy brain (Table S3; Fig. S1A,B). However, GSEA of cortistatin-deficient ischemic brain samples revealed significant upregulation of innate immunity, antiviral defence, pattern recognition receptor signalling, pro-inflammatory cytokine responses (IFN-γ, IL-6, and TNF), and extracellular matrix (ECM) remodelling (Table S4; Fig. S1C).

Next, to identify cortistatin-specific responses to ischemia, we compared transcriptomic data from contralateral to ischemic hemispheres within each genotype (Fig. 2A). *Cort*^+/+^ mice exhibited 443 DEGs (419 upregulated and 24 downregulated in the injured tissue), whereas *Cort*^−/−^ mice showed a markedly heightened response with 1483 DEGs (1213 upregulated and 270 downregulated) (Fig. 2B; Tables S5, S6). Although substantial DEG overlap was observed between genotypes (Fig. 2C), *Cort*^−/−^ mice demonstrated a more robust enrichment across biological processes (BP) terms including innate and adaptive immune activation, leukocyte infiltration, inflammation, cell cycle progression, ERK1/ERK2, JAK-STAT and MAPK signalling, neuroinflammation, gliogenesis, ECM remodelling, angiogenesis, and ROS metabolism (Fig. 2D; Table S7). Additional enrichment was observed in cellular components (CC) related to membrane specialization, ECM structure; mitotic spindle formation, and adhesion-related protein complexes (Table S8; Fig. S2); as well as in molecular functions (MF) such as immune receptor activity, integrin and molecular factor binding, cytoskeletal remodelling, and GTPase and kinase activity (Table S9; Fig. S3). Importantly, we validated the expression of key upregulated and downregulated genes by qPCR, showing a high degree of correlation with the RNA-seq quantification (Fig. S4).

**Figure 2.**
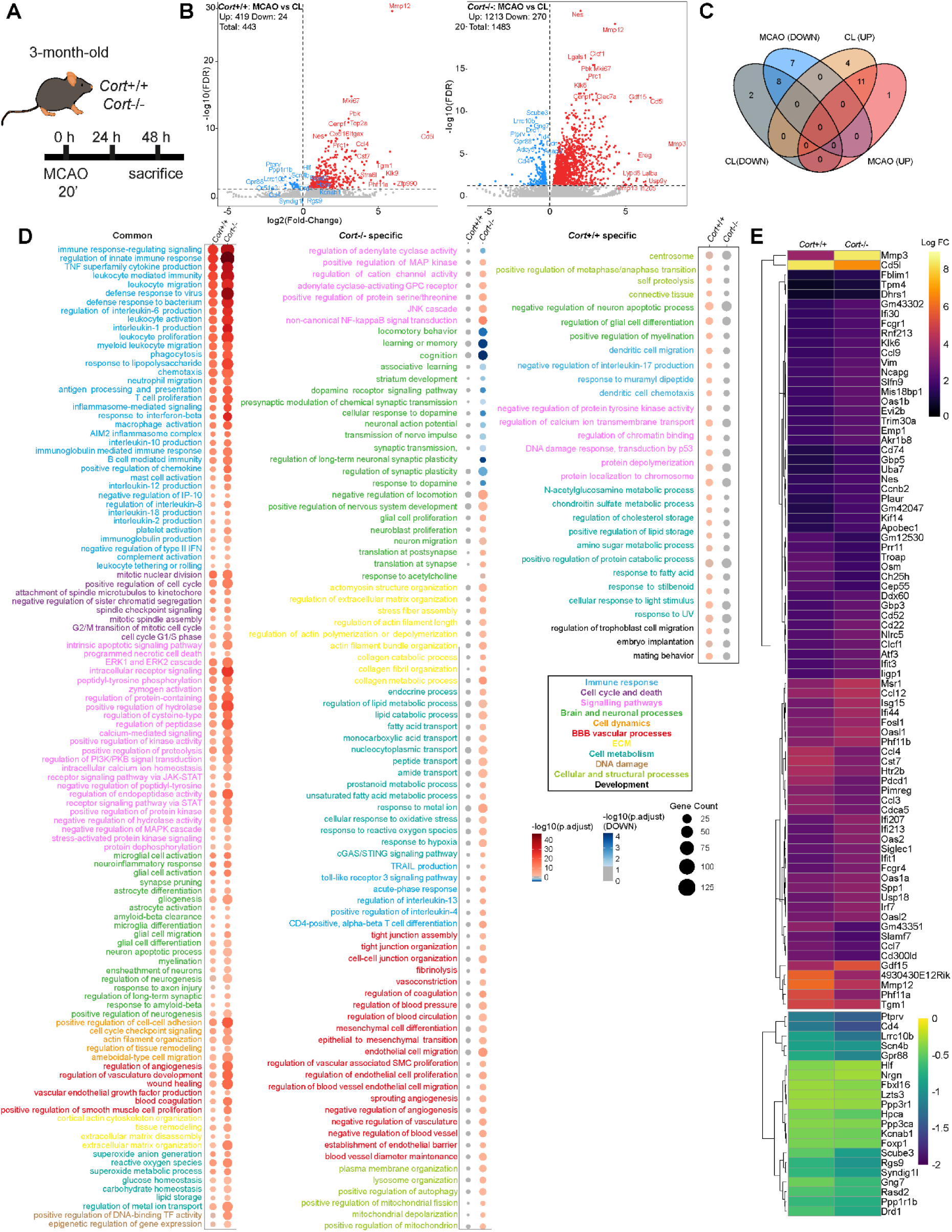
Lack of cortistatin leads to an exaggerated transcriptional response to cerebral ischemia. **(A)** Transcriptomic profiling was performed on *Cort^−/−^* and *Cort^+/+^* mouse brains isolated 48 h after ischemic stroke induction (20 min MCAO). **(B)** Volcano plots depict differentially expressed genes (DEGs) in ischemic (MCAO) *vs.* healthy contralateral (CL) hemispheres from *Cort^+/+^* (left) and *Cort^−/−^*(right) mice. Each dot represents one gene. Grey dots represent non-significantly altered genes. Significantly upregulated (red) and downregulated (blue) genes (with false discovery rate, FDR <0.05) during ischemic injury are shown. Comprehensive DEG lists are available in Additional file 1: Tables S5 and S6. **(C)** The Venn diagram displays the overlap of significantly upregulated and downregulated genes in *Cort^−/−^* and *Cort^+/+^* mice. **(D)** Gene ontology (GO) enrichment analysis of biological processes revealed distinct and shared terms between healthy and ischemic brain in *Cort^−/−^*compared to *Cort^+/+^*. DEGs were manually categorized into functional networks including: immune response (blue), cell cycle and death (purple), signalling pathways (pink), brain and neuronal processes (green), cell dynamics (orange), BBB and vascular processes (red), extracellular matrix (ECM, yellow), cell metabolism (turquoise), DNA damage (brown), cellular and structural processes (pistachio green), and development (black). Shared and genotype-specific terms are indicated. All GO terms and corresponding genes are listed in Additional file 1: Table S7. Colour intensity represents statistical significance (red: upregulated, blue: downregulated, grey: non-significant). Circle size reflects the number of DEGs in each pathway. **(E)** Heatmap of Log_2_FC values for common DEGs across both genotypes. Hierarchical clustering is shown for shared upregulated (top, selected 78/407 genes) and downregulated (bottom, 21/21 genes) transcripts. Full description of DEGs provided in Additional file 1: Table S10.

Beyond shared responses to ischemia, cortistatin-deficient mice exhibited distinct upregulation of BP involving JNK and non-canonical NF-kappaB signalling, glial proliferation, ECM organization, lipid and amide metabolism, BBB regulation, and mitochondrial integrity. Unique enrichment in CC and MF categories included endosomal/lysosomal compartments, MHC complexes, cell division, cytoskeletal components, Golgi and ribosomal membranes (CC); and inhibitor and protein kinase activities, transcriptional regulation, molecular binding, and transporter function (MF). In contrast, terms exclusively enriched in *Cort*^+/+^ animals primarily involved regulation of glial differentiation, myelination, lipid/cholesterol storage, and mating behaviour (BP); cell cycle and Ser/Thr complexes (CC); oxidoreductase activity, hyaluronic and ferric iron binding (MF). Strikingly, only *Cort*^−/−^ mice demonstrated significant downregulation of gene sets associated with cognition, motor behaviour, dopaminergic signalling, synaptic plasticity (BP); neuronal spine, synapses, and postsynaptic structures (CC); and channel, neurotransmitter and glutamate receptor activities (MF) (Fig. 2D; Tables S7-9; Fig. S2,3).

We found 428 DEGs shared between wild-type and cortistatin-deficient mice (407 upregulated, 21 downregulated in ischemia; Table S10). A more detailed analysis revealed genotype-specific expression changes when comparing Log_2_FC differences (Log_2_FC *Cort*^+/+^ (MCAO *vs.* CL) – Log_2_FC *Cort*^−/−^ (MCAO *vs*. CL)) (Fig. 2E). Genes with greater upregulation in *Cort*^−/−^ animals were associated with immune activation, persistent neuroinflammation, gliosis, BBB breakdown, ECM and vascular remodelling, oxidative stress, and impaired neuroprotection (Table S11). Additionally, several genes with undefined roles in stroke but highly upregulated in *Cort*^−/−^ mice may contribute to glial priming and post-ischemic inflammation. Conversely, *Cort*^+/+^ mice showed enhanced upregulation of genes involved in inflammation resolution, tissue repair, and neurogenesis (Table S12). Genes more strongly downregulated in *Cort*^−/−^ mice, though their roles in stroke are not fully established, have been implicated in neuroinflammation, vascular instability, impaired synaptic plasticity, disrupted GPCR, and dopamine signalling in other contexts (Table S13).

These findings demonstrate that, independently of cortistatin, a common molecular signature is displayed as a general response to ischemia. However, genetic ablation of cortistatin amplifies this transcriptional response, leading to exacerbated neuroinflammation and compromised neurovascular remodelling, and leads to unique pathways characterized by the repression of responses critical for synaptic function and recovery.

### Ischemic stroke induces the downregulation of both cortistatin and its receptors

To understand the mechanistic basis of this vulnerability, we evaluated the availability of cortistatin-binding receptors (*Sstr1–5* and *Ghsr*). All receptors were expressed in the brain, with *Sstr4* and *Sstr2* showing the highest relative levels (Fig. S5). However, ischemia induced a marked and widespread reduction in their expression in the ipsilateral hemisphere across both genotypes. Notably, although baseline levels in the contralateral hemisphere were comparable, stroke-induced receptor downregulation was more pronounced in cortistatin-deficient mice (Fig. S6). In parallel, cortistatin expression itself was strongly reduced following stroke (Fig. 1).

### Lack of cortistatin exacerbates cell-type marker gene profiles following ischemic stroke

To elucidate genotype-dependent cellular responses to cerebral ischemia, we employed computational deconvolution of bulk transcriptomic data to perform a predictive Marker Gene Profile (MGP) analysis, estimating the relative enrichment of cell-type-specific markers in the presence or absence of endogenous cortistatin. By using MGPs, we estimated relative shifts in specific transcriptional signatures of neuronal, vascular, immune, and glial cell types (Fig. S6; Table S14). While this well-established bioinformatic approach does not substitute for absolute single-cell quantification, it provides a robust estimation of overarching population-level dynamics. Among neuronal populations, both genotypes exhibited a significant reduction in spiny neuron signatures following MCAO, consistent with established features of ischemic injury [52]. Notably, a selective decrease in gene markers of dentate granule neurons was observed exclusively in cortistatin-deficient mice, indicating increased vulnerability of this neuronal subtype (Fig. S6A). Ependymal neuron marker abundance increased in both genotypes post-ischemia, more prominently in *Cort^−/−^* mice, potentially reflecting a transcriptomic signal compensatory neurogenesis. Interestingly, the signatures for GABAergic interneuron populations (as cortistatin-expressing neurons), dopaminergic and NPCs remained unchanged following stroke (Fig. S6A). Additionally, MGP analysis revealed that only *Cort^−/−^* mice had significant increase in gene expression signatures of endothelial cells, pericytes, and vascular smooth muscle cells, suggesting an amplified transcriptomic vascular response in the absence of cortistatin (Fig. S6A). This was corroborated by GSEA enrichment (Fig. S6B). Both approaches also demonstrated that signatures of CNS-associated macrophages, lymphocytes, monocyte-derived cells, and neutrophils were enriched in the ischemic hemisphere of both genotypes, but were significant only in *Cort*^−/−^ mice, indicating heightened immune cell-marker footprint (Fig. S6).

Analysis of glial cell populations markers demonstrated a significant increase in both activated and deactivated microglial gene signatures in *Cort^−/−^* mice, which, together with peripheral immune marker subsets, indicate a dysregulated neuroimmune response post-stroke. Astrocyte– and oligodendrocyte-specific markers also revealed divergent phenotypes: a “Type 1” signature profile (based on markers from [43]) was significantly reduced in *Cort^−/−^*mice following MCAO, whereas a “Type 2” profile (characterized by markers from [42], was significantly increased) (Fig. S6A; Table S14).

Downregulation of astrocytic “Type 1” markers, typically associated with homeostatic and neuroprotective functions, suggests a shift toward reactive or dysfunctional astrocyte signatures in cortistatin-deficient mice following ischemia. Concurrent upregulation of the “Type 2” profile indicates the emergence of a heterogeneous reactive astrocyte population encompassing both neurotoxic and neuroprotective subsets. Although not statistically significant, the oligodendrocyte profiles exhibited similar opposing trends in both genotypes: decreased “Type 1” and increased “Type 2” signatures (Fig. S6A). Analysing the markers selective for each cell population (Table S14), the elevated “Type 2” profile likely reflects an injury-induced remyelination response, whereas the greater reduction of “Type 1” markers in *Cort^−/−^*mice may denote impaired oligodendrocyte maturation and myelin maintenance.

These findings underscore the critical role of cortistatin in preserving cellular homeostasis post-stroke. Its deficiency drives broad dysregulation across neuronal, vascular, immune, and glial compartments, fostering a pathological neuroimmune and neurovascular environment that, together with an altered receptor network, may hinder tissue repair and functional recovery.

### Cortistatin deficiency exacerbates stroke-induced neurodegeneration, which is mitigated after cortistatin treatment

To investigate the functional significance of the differential molecular pathways driven by endogenous cortistatin in stroke pathophysiology, we subjected *Cort*^−/−^ mice and age-matched wild-type mice to mild MCAO and assessed acute neuropathological outcomes.

Our findings showed that cortistatin-deficient mice exhibited a greater propensity for extensive infarcts, with 37.5% displaying lesions >30 mm^3^, while all *Cort*^+/+^ animals presented smaller lesions, as determined by CV staining (Fig. 3A,B; Table S15). MAP-2 immunostaining confirmed significant pronounced neuronal injury in *Cort^−/−^* mice, with >5 mm^2^ MAP-2-negative areas in 75% of animals, compared to only 12.5% wild-type controls (Fig. 3A,B; Table S15). Furthermore, neurological impairment was significantly aggravated in *Cort^−/−^* mice, with 30% scoring 2-3 (persistent circling, severe motor incoordination, tremors), whereas most of wild-type mice remained below a score of 2 (intermittent circling, moderate motor incoordination, and mild hypomobility) (Fig. 3C).

**Figure 3.**
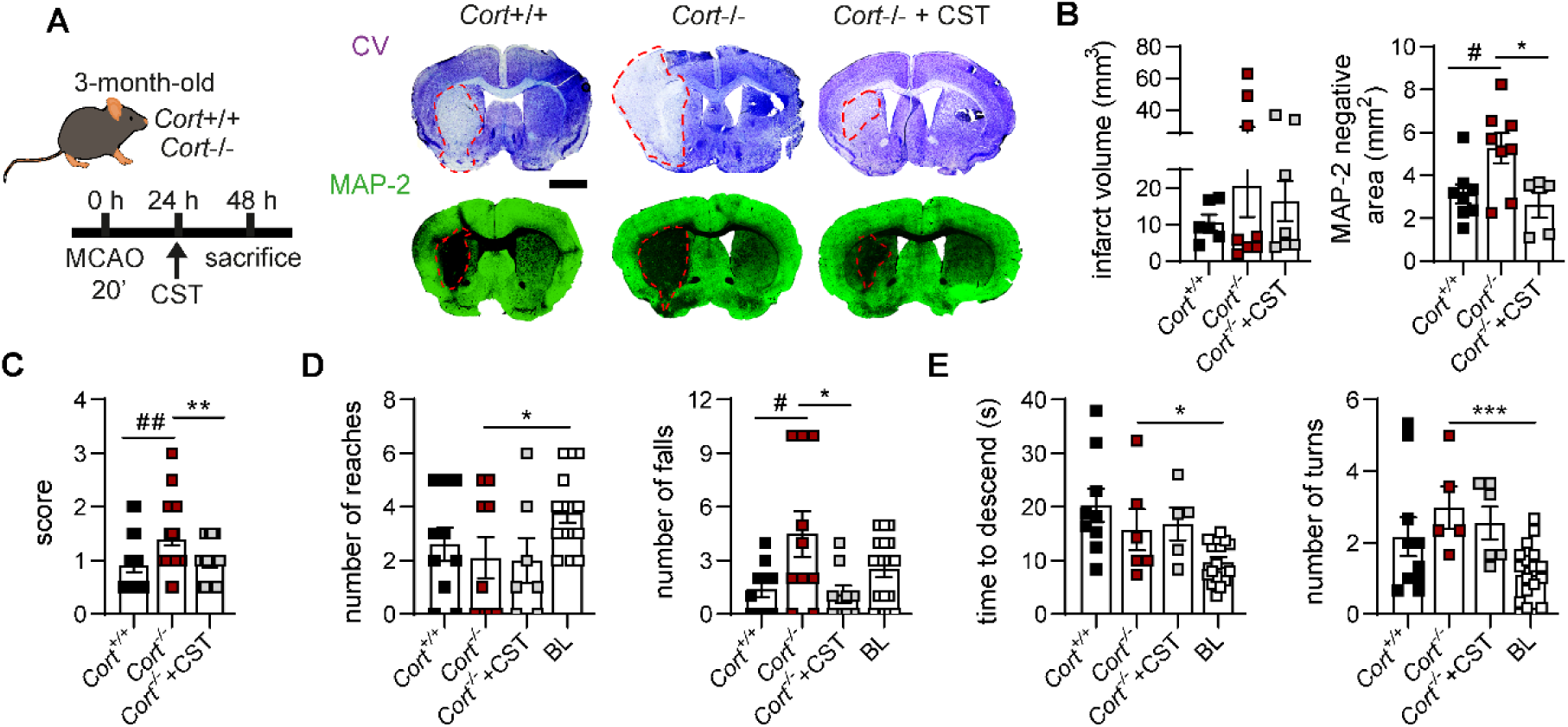
Cortistatin deficiency exacerbates neuronal injury and functional outcomes. **(A)** Left, *Cort*^−/−^ mice underwent MCAO and received saline or cortistatin (144μg/kg) 24 h after reperfusion and were sacrificed 48 h post-surgery. Animals were compared with *Cort*^+/+^ MCAO saline-treated mice. Right, coronal brain sections were stained with cresyl violet (CV) (upper panel) or MAP-2 (lower panel, green) to identify neuronal damage. Ischemic lesions are delineated by dashed red lines (pale/white area in cresyl violet and loss of immunoreactivity/dark area in MAP-2). Scale bar: 1,500 µm. **(B)** Infarct volume quantified from 10 CV-stained brain sections (0.5 mm intervals, left) or by the absence of MAP-2 on coronal sections (right). N=5-9 mice/group. **(C)** Neurological scores evaluated at 48 h post-MCAO (0, no signs; 3, death). N=17-29 mice/group. **(D-E)** Motor performance in wire-hanging (D) and pole (E) tests, at baseline (BL, pre-MCAO) and 48 h post-MCAO. **(D)** Number of reaches towards one of the posts (left) and falls (right). N=7-10 mice/group. **(E)** Time to descend (left) and number of turns (right) until reaching the pad. N=5-6 mice/group. Data represent mean ± SEM. Dots indicate individual mice. Statistical analyses were performed using unpaired t-tests or one-way ANOVA for comparisons among multiple groups. \**vs*. *Cort*^−/−^ MCAO saline-treated mice, ^#^*vs*. *Cort*^+/+^ MCAO saline-treated mice, *^/#^*p*≤0.05, **^/#/#^*p*≤0.01, ****^/###/#^*p*≤0.0001 CST: cortistatin

Functional deficits in cortistatin-deficient mice after stroke were further evidenced by impaired performance in wire-hanging and pole tests compared to baseline (Fig. 3D,E). Although motor deficits in the behavioural test were comparable between genotypes, cortistatin-deficient mice displayed significantly more frequent falls (40% fell >5 times *vs.* none in wild-type). Cortistatin-deficiency also seems to affect the success rate of reaching the post in the wire-hanging test (*i.e.,* over 40% of cortistatin-deficient animals never reached the post, whereas 80% of wild-type animals reached it at least once) (Fig. 3D; Table S16).

Remarkably, exogenous cortistatin administration to *Cort*^−/−^ mice following MCAO significantly ameliorated neuronal injury (*i.e.,* over 70% of treated animals exhibited a lesion volume <30 mm^3^ and none showed MAP-2-negative areas >5 mm^2^). It also improved neurological score and restored some motor outcomes to levels comparable with wild-type controls (Fig. 3A-D; Tables S15, S16).

These results underscore the heightened vulnerability to ischemic stroke in the absence of endogenous cortistatin and highlight its therapeutic role in mitigating stroke-induced damage.

### The neuroimmune response after stroke is dysregulated in cortistatin-deficient mice but becomes neuroprotective following cortistatin administration

The neuroimmune response to stroke involves intricate and coordinated CNS-peripheral immune interactions that shape both acute injury and long-term brain recovery [53]. Our transcriptomic data revealed sustained neuroinflammation in cortistatin-deficient mice post-stroke. However, the intrinsic role of cortistatin in modulating systemic immune responses after stroke remains unexplored. In this context, we found that, compared to wild-type mice, animals lacking cortistatin exhibited significantly elevated IL-6 and MCP-1 serum levels post-MCAO, accompanied by reduced spleen mass and a trend towards lymphopenia, hallmarks of dysregulated peripheral immunity (Fig. 4A,B). Administration of cortistatin normalized IL-6, elevated IL-12, restored spleen weight and increased splenocyte count, indicating reestablishment of systemic immune balance (Fig. 4A,B).

**Figure 4.**
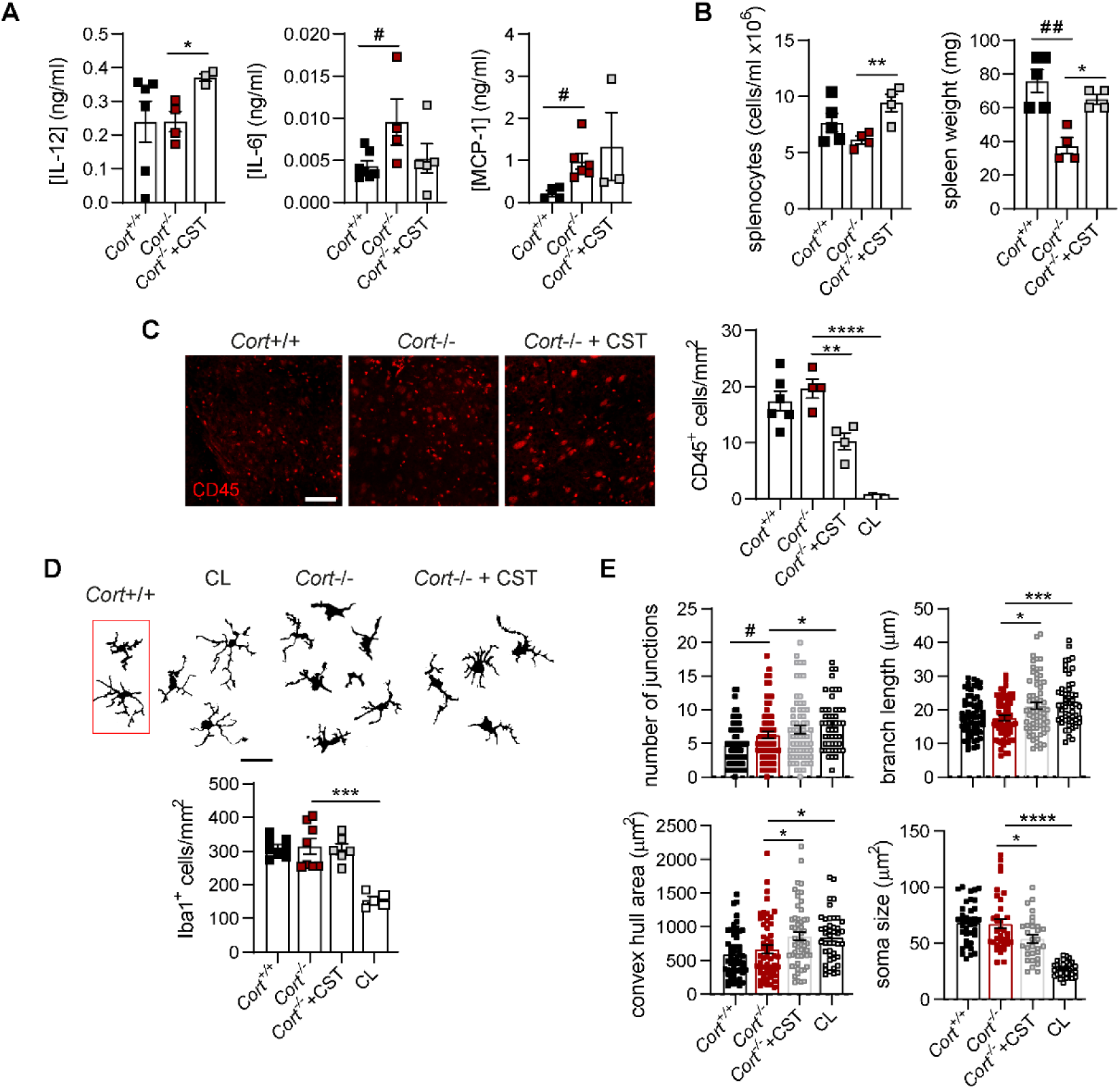
Lack of cortistatin triggers a dysregulated immune response. *Cort*^−/−^ mice underwent MCAO and received saline or cortistatin (144μg/kg) 24 h after reperfusion and were sacrificed 48 h post-surgery. Animals were compared with *Cort*^+/+^ MCAO saline-treated mice. **(A)** Serum levels of IL-12, IL-6, and MCP-1 (ng/ml) measured by ELISA. IL-12: N=3-6; IL-6: N=4-7; MCP-1: N=3-6. **(B)** Spleen cellularity and weight (mg). N=4-5. **(C)** Left, CD45-stained coronal sections showing leukocytes (red). Scale bar: 100 μm. Right, leukocyte density (cells/mm^2^) in the ischemic striatum N=3-6. **(D)** Top, representative binarized microglia from contralateral (CL) or peri-lesion regions. Red box: *Cort*^+/+^ MCAO saline-treated. Scale bar: 30 μm. Bottom, Iba1+ population (cells/mm^2^) in the CL and ischemic core regions N=5-8. **(E)** Iba1^+^ cells morphology assessed using skeleton (junctions, branch length; n=50-65 cells, N=3 mice/group) and fractal (convex hull area; n=43-55 cells, N=3 mice/group) analyses. Soma size was manually measured (n=30-35 cells, N=3 mice/group). Data represent mean ± SEM. Statistical analyses were performed using unpaired t-tests or one-way ANOVA for comparisons among multiple groups. \**vs*. *Cort*^−/−^ MCAO saline-treated mice, ^#^*vs*. *Cort*^+/+^ MCAO saline-treated mice, *^/#^*p*≤0.05, **^/#/#^*p*≤0.01, ****^/###/#^*p*≤0.0001 CST: cortistatin

In the brain, elevated densities of CD45^+^ and Iba1^+^ myeloid cells were observed in the infarct core of cortistatin-deficient mice, paralleling wild-type levels (Fig. 4C,D). However, cortistatin deficiency exacerbated post-stroke myeloid cell activation. Compared to contralateral hemisphere, Iba1^+^ cells from mice lacking cortistatin displayed a reactive morphology, characterized by reduced branching, shorter processes, increased soma size, and diminished territory area, suggesting enhanced activation (Fig. 4D,E). Interestingly, branching was significantly higher than in wild-type mice. Gene expression analysis revealed similar upregulation of both pro– and anti-inflammatory markers between genotypes (*Trem2, CD68, Iba1, TNF-α*, *IL-6* and *Arg1*), though *Cort^−/−^* mice showed significantly reduced *CD163* and *IL-10* expression, indicative of impaired anti-inflammatory responses (Fig. S7). Cortistatin administration to cortistatin-deficient mice selectively reduced CD45^+^ infiltrating populations without affecting Iba1^+^ cells, suggesting distinct regulatory mechanisms. Indeed, cortistatin treatment promoted a shift toward a more ramified Iba1^+^ phenotype with increased IL-10, supporting a reparative glial response distinct from MCAO wild-type profiles (Fig. 4C-E; Fig. S7B). Consistent with these results, the anti-inflammatory action of cortistatin was further demonstrated in primary adult microglia, astrocytes, and mixed neuronal-glial cultures subjected to inflammatory (LPS) or ischemia-like conditions (OGD+LPS). Under these conditions, cortistatin-deficient cells exhibited an exacerbated pro-inflammatory phenotype (*e.g.,* elevated levels of TNF-α, IL-6, and NO) that was significantly attenuated by exogenous cortistatin treatment *(i.e.,* reducing cytokine levels) (Fig. S8).

Together, these findings demonstrate that cortistatin is essential for maintaining neuroimmune and glial homeostasis after stroke. Cortistatin deficiency drives excessive peripheral and central immune activation, coupled with dysfunctional glial responses, thereby amplifying neuronal injury. Conversely, exogenous cortistatin reprograms both systemic and CNS immune compartments toward reparative phenotypes.

### Cortistatin deficiency affects BBB integrity and vascular dynamics after ischemic stroke, which are normalized following cortistatin administration

Next, we investigated BBB dynamics post-stroke. Following MCAO, cortistatin-deficient mice exhibited significantly elevated parenchymal IgG accumulation relative to wild-type controls, indicative of substantial BBB disruption (Fig. 5A). This permeability likely contributes to the enhanced immune cell infiltration previously observed (Fig. 4C). Analysis of vascular markers revealed that C*laudin-1*, *Occludin*, and *Col1a2* levels were significantly reduced in the ischemic hemisphere of cortistatin-deficient mice compared to the contralateral side, while C*laudin-5* expression remained unchanged (Fig. 5B). Notably, these TJ expression levels were similar to the ones found in wild-type mice. However, ZO-1 distribution was significantly disrupted in the absence of cortistatin, with notable loss across the endothelium (Fig. 5C). A comparable vessel density and diameter post-MCAO were observed between genotypes (Fig. 5D). Administration of cortistatin to cortistatin-deficient mice significantly reduced IgG leakage, normalised *Claudin-1* and *Occludin* expression, and restored ZO-1 configuration, thereby attenuating BBB disruption (Fig. 5A-C). In addition, cortistatin treatment promoted vascular remodelling, evidenced by increased vessel area and diameter within the ischemic striatum (Fig. 5D). Although TJ architecture and endothelial integrity were similarly compromised between genotypes, we observed that astrocytic responses, intrinsically related with neuroinflammation, vascular remodelling and BBB function, were disrupted in cortistatin-deficient mice. In the absence of cortistatin we found reduced glial scar formation and atypical astrocyte morphologies compared to wild-type animals (smaller soma, fewer and shorter processes and elongated morphology, increased span ratio, and reduced territorial coverage) (Fig. 5E,F). Additionally, GFAP expression, upregulated after MCAO, was significantly lower in cortistatin-deficient mice compared to wild-type animals. This was accompanied by a decrease in BDNF and GDNF compared to the contralateral hemisphere (Fig. 5G). Cortistatin treatment markedly restored astrocytic morphology and function and increased GDNF expression to similar levels seen in wild-type MCAO mice (Fig. 5E-G).

**Figure 5.**
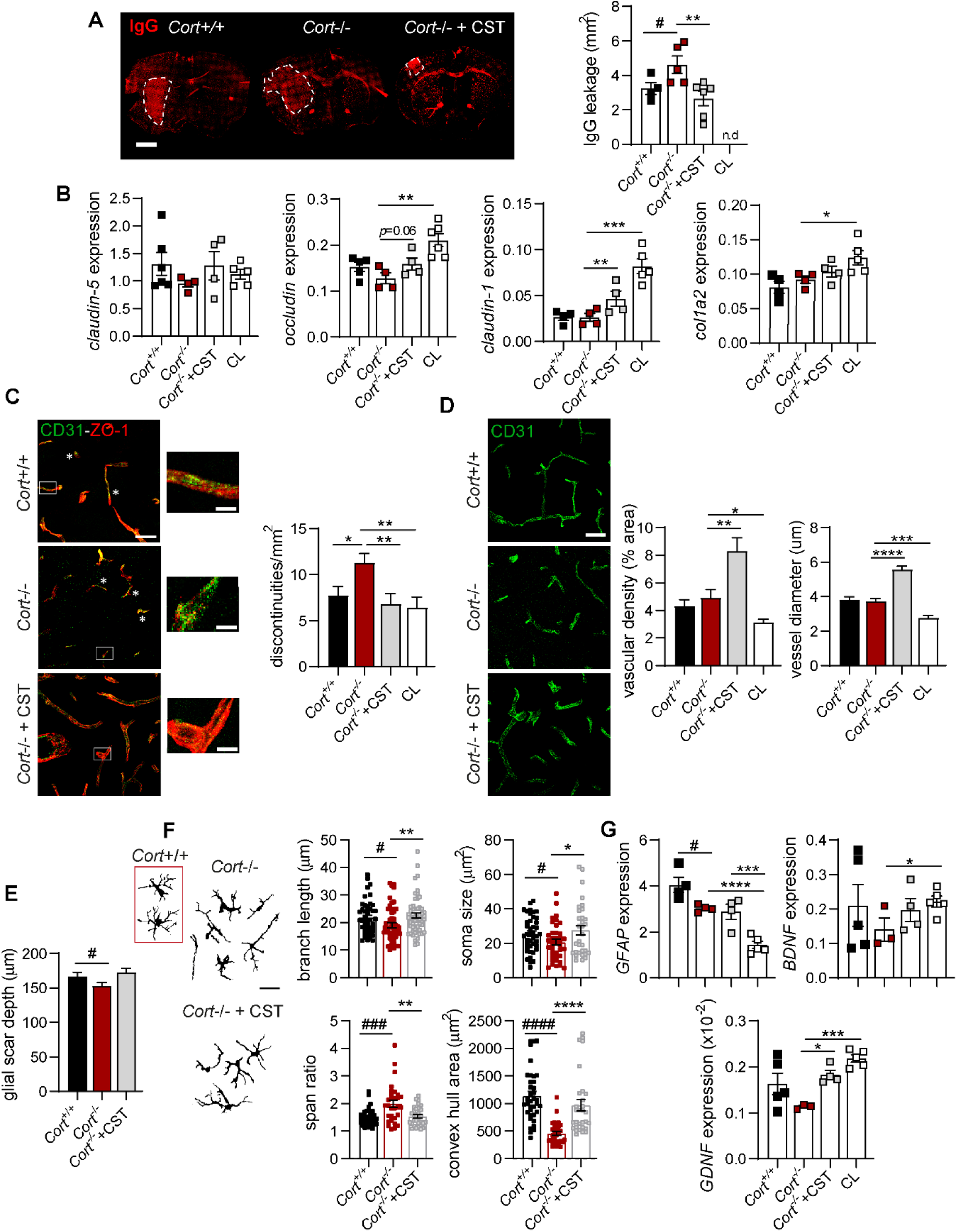
Cortistatin loss disrupts astrocytic function and BBB integrity post-stroke. MCAO was conducted in *Cort*^−/−^ mice treated with saline or cortistatin (144 μg/kg) 24 h after reperfusion, and sacrificed 48 h later. Mice were compared to *Cort*^+/+^ MCAO saline-treated mice. **(A)** Left, coronal brain sections stained with fluorescently labelled anti-mouse IgG show leakage areas (dashed white lines). Scale bar: 1,500μm. Right, quantification of IgG^+^ areas. N=5-6 mice/group. **(B)** Gene expression levels of C*laudin-5, Claudin-1, Occludin*, and *Col1a2* in contralateral (CL) and ipsilateral brain hemispheres normalized to *Rplp0*. N=4-6 mice/group. **(C)** Left, representative images of the ipsilateral striatum from *Cort*^−/−^ mice post-MCAO and treated with cortistatin (CST), stained for CD31 (brain endothelial cells, green) and ZO-1 (TJs, red). Scale bar: 30μm. Insets highlight ZO-1 discontinuities (asterisks) and recovery of ZO-1 coverage (arrows). Scale bar: 120μm. Right, quantification of ZO-1 discontinuities per vessel area. n=2 ROIs/mouse; N=3-6 mice/group. **(D)** Left, CD31 staining in the ipsilateral striatum of *Cort*^−/−^ mice (green). Scale bar: 30μm. Right, vascular density (CD31-positive area/total image area; n=3 ROIs/mouse; N=3-5 mice/group) and vessel diameter were quantified in different sections from each vessel (5-10 vessels/mouse, N=3-4 mice/group). **(E)** Glial scar thickness was quantified. n=54-57 ROIs; N=4 mice/group. **(F)** Left, binary images of astrocytes from the glial scar. Scale bar: 30μm. Right, quantification of astrocyte morphology, including branch length (skeleton analysis, n=49–57 cells), soma size (manual, n=32–39 cells), span ratio, and convex hull area (fractal analysis, n=30 cells; N = 3 mice/group). **(G)** Gene expression levels of *GFAP*, *BDNF,* and *GDNF* in CL and ipsilateral hemispheres of *Cort*^−/−^ mice treated with saline or cortistatin following MCAO, normalized to *Rplp0*. N=3-5 mice/group. Data represent the mean ± SEM. Statistical analyses were performed using unpaired t-tests or one-way ANOVA for comparisons among multiple groups. \**vs. Cort*^−/−^ MCAO saline-treated mice, ^#^*vs. Cort*^+/+^ MCAO saline-treated mice, *^/#^*p*≤0.05, **^/#/#^*p*≤0.01, ***^/##/#^*p*≤0.001, ****^/####^*p*≤0.0001.

These results indicate that cortistatin deficiency impairs astroglial scar formation and exacerbates BBB breakdown and vascular instability, while exogenous cortistatin restores glial support, barrier integrity and vascular resilience.

### Absence of cortistatin impacts subacute stroke outcome while exogenous treatment potentiates recovery

Regarding the role of cortistatin in acute stroke, and following STAIR recommendations [54,55], we extended our analysis to the early long-term post-stroke stage (7 days post-MCAO), focusing on key markers of injury and recovery. While neurological deficit scores were initially comparable between genotypes (Fig. 6A), animals lacking cortistatin exhibited trends toward larger infarcts, greater MAP-2-negative areas, and reduced neuronal density relative to wild-type controls (Fig. 6B; Table S15; Fig. S9A). Functional impairments persisted in *Cort*^−/−^ mice, including decreased successful reaches and poorer wire-hanging test performance, although pole test results remained unaffected (Table S16; Fig. S10A,B). *Cort^−/−^* mice also demonstrated cognitive impairments in the novel object recognition test, exhibiting a significantly lower discrimination and recognition index at baseline compared to wild-type animals (Fig. S11). Following the ischemic insult, although no significant differences were observed between genotypes, the cortistatin-treated groups showed a positive trend toward recovery, indicating an amelioration of this cognitive deficit. Importantly, histological analysis revealed pronounced axonal disorganization and aberrant myelin structures in *Cort*^−/−^ mice, reflected by elevated MBP intensity in the ischemic hemisphere, suggestive of myelin disruption (Fig. 6C). Cortistatin treatment ameliorated these pathological alterations, improving neuronal density, axonal integrity, and neurological scores in both genotypes (Fig. 6A-C).

**Figure 6.**
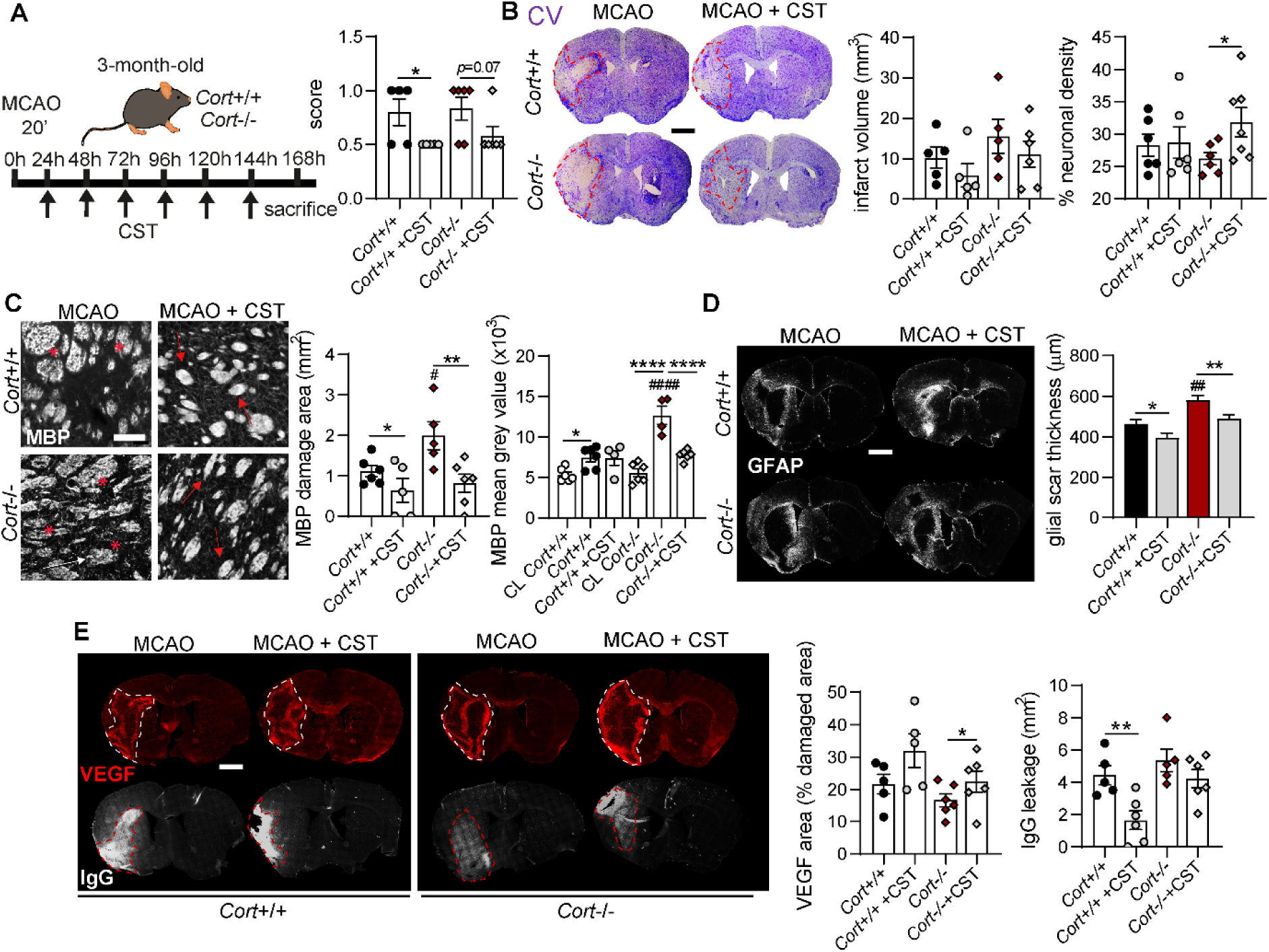
Cortistatin deficiency impairs long-term stroke recovery, while treatment improves subacute outcomes. **(A)** *Cort^+^*^/+^ and *Cort*^−/−^ MCAO mice were treated daily with either saline or cortistatin (144μg/kg), from 24 h post-reperfusion, and sacrificed at 7 days (left). Neurological scores were evaluated at 7 days using a scale ranging from 0 (no signs) to 3 (death) (right). N=5-6 mice/group. **(B)** Left, brain coronal sections were stained with cresyl violet (CV) to assess neuronal damage (pale/white area). Ischemic lesions are outlined with dashed red lines. Scale bar: 1,500 µm. Centre, ischemic volume was determined by analysing ten CV-stained brain slices at 0.5 mm intervals per mouse. Right, neuronal density within the ischemic area was quantified (% of total area). N=4-6 mice/group. **(C)** Left, representative images of the ipsilateral striatum stained with MBP antibody to visualize myelin sheaths. Aberrant myelin sheaths are depicted with asterisks while recovered ones are marked with arrows. Scale bar: 30 μm. Right, the area of damaged myelin sheaths (left) and MBP mean grey value (right) were quantified. N=4-6 mice/group. **(D)** Left, representative brain sections stained with GFAP antibody to identify astrocytes. Scale bar: 1,500 μm. Right, glial scar thickness was quantified. N=4-6 mice/group. **(E)** Left, sections were stained with VEGF antibody (red) and with a fluorescently-labelled antibody against mouse IgGs (white). Scale bar: 1,500 μm. Centre, VEGF^+^ area was quantified as the % of positive area within the damaged area. N=4-6 mice/group. Right, IgG^+^ area was measured to assess BBB leakage. N=4-6 mice/group. Data represent the mean ± SEM. Dots indicate individual mice. Statistical analyses were performed using unpaired t-tests or one-way ANOVA for comparisons among multiple groups. \**vs. Cort^+/+^* or *Cort^−/−^* MCAO saline-treated mice, ^#^*vs. Cort^+^*^/+^ MCAO saline-treated mice, *^/#^*p* ≤ 0.05, **^/#/#^*p* ≤ 0.01, ****^/####^*p* ≤ 0.0001

Glial responses were also altered in the subacute phase. Iba1+ cell density increased substantially compared to acute timepoints (>10^3^ cells/mm^2^ *vs.* <400 cells/mm^2^) with no genotype– or treatment-dependent differences (Fig. S9B). Similarly, glial scar thickness augmented as the stroke progressed (<200 to >400 µm from acute to subacute stroke), with *Cort*^−/−^ mice transitioning from disorganized to significantly thicker fibrotic scars at 7 days (Fig. 5E, 6D). Cortistatin treatment reduced scar thickness in both genotypes, suggesting a role in regulating scar maturation. Additionally, cortistatin administration to cortistatin-deficient mice upregulated VEGF expression in the ischemic hemisphere, potentially enhancing angiogenesis and BBB repair (Fig. 6E). Consistently, IgG extravasation was reduced following treatment in wild-type animals, aligning with observations at earlier stages (Fig. 5A, Fig. 6E).

These findings demonstrated that cortistatin deficiency worsens early long-term stroke pathology, hindering neurovascular and glial repair, and according to transcriptomic studies, showing limited activation of repairing pathways in the ischemic brain without cortistatin. Exogenous cortistatin reversed many of these pathological features, underscoring its broad neurorepair potential beyond the acute phase.

### Cortistatin regulates neuroinflammatory and vascular outcomes after stroke in 6-month-old mice

Next, we evaluated the influence of cortistatin on stroke outcome in the context of premature aging, by inducing a mild MCAO in 6-month-old *Cort*^+/+^ and *Cort*^−/−^ mice, with or without cortistatin treatment, and compared results to 3-month-old MCAO animals. While standard aging models typically utilize 12–18-month-old animals, we specifically selected this 6-month time point to investigate the accelerated aging phenotype inherent to cortistatin-deficient mice, as previously described by our laboratory [56]. Crucially, this age marks the upper limit for feasible surgical stroke induction, as older cortistatin-deficient mice show very high mortality after MCAO. This age group also aligns with reported declines in functional performance in C57BL/6J males [57].

In the acute phase, cortistatin deficiency in 6-month-old mice led to worsened neurological scores and behavioural deficits, paralleling findings in younger cohorts (Fig. 7A; Fig. S10C,D). Neuronal damage, as indicated by MAP-2 loss, was significantly higher in older wild-type mice compared to their younger counterparts, a difference not observed between 3-month-old and 6-month-old *Cort*^−/−^ animals (Fig. 7B; Table S15). Cortistatin treatment reduced neuronal injury in wild-type mice and led to an improved score in cortistatin-deficient mice (Fig. 7A,B). Additionally, cortistatin treatment improved some behavioural outcomes in the pole test, though it demonstrated generally limited effect (Table S16; Fig. S10C,D).

**Figure 7.**
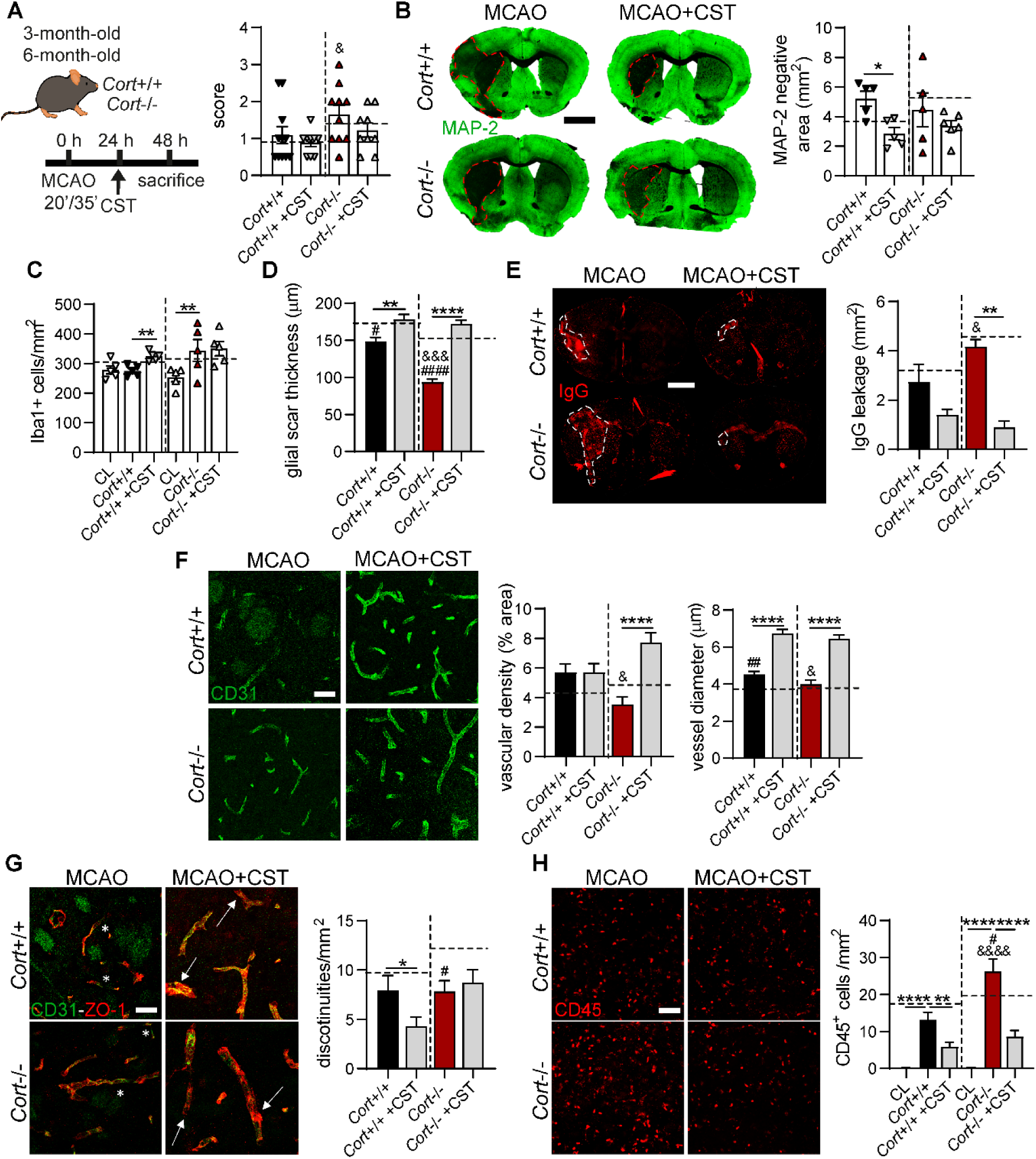
Lack of cortistatin in 6-month-old mice worsens stroke outcomes, promoting glial dysregulation and vascular disruption. **(A)** 6-month-old *Cort^+/+^* and *Cort^−/−^* mice were subjected to MCAO and treated with saline or cortistatin (144μg/kg) 24 h post-reperfusion and sacrificed at 48 h (left). Neurological deficits were scored from 0 (no signs) to 3 (death) (right). N=9-12 mice/group. **(B)** Left, MAP-2 staining. Ischemic areas (loss of immunoreactivity/dark area) are delineated with red dashed lines. Scale bar: 1,500 μm. Right, quantification of MAP-2 loss. N=5–6 mice per group. **(C)** Iba1^+^ population (cells/mm²) quantified in contralateral (CL) and ischemic core regions. **(D)** Glial scar thickness was quantified, n=30-35 ROIs, N=3 mice/group. **(E)** Left, brain sections stained with fluorescently labelled anti-mouse IgG show leakage areas (dashed white lines). Scale bar: 1,500 μm. Right, IgG^+^ area was quantified to assess BBB permeability. N=4–6 mice/group. **(F)** Left, images of the lesion core (striatum) stained with CD31 (green). Scale bar: 30μm. Vascular density (CD31^+^ area/total area; n=3 ROIs/mouse; N=3-4 mice/group) and vessel diameter quantified in sections from each vessel (5-10 vessels/mouse, N=3-6 mice/group). **(G)** Left, images of the ipsilateral striatum stained with CD31 (green) and ZO-1 (red) to assess TJ integrity. ZO-1 discontinuities (asterisks), and ZO-1 uniformity (arrows). Scale bar: 30 μm. Right, quantification of ZO-1 discontinuities per vessel area. n=3 ROIs/section/mouse; N=3–5 mice/group. **(H)** Left, high-magnification images (striatum) stained with CD45 (red) to detect infiltrating leukocytes. Scale bar: 100 μm. Right, leukocyte density (cells/mm²) quantified in the ischemic striatum and the CL hemisphere. All data were compared to 3-month-old MCAO *Cort^+/+^* and *Cort^−/−^* mice treated with saline (dashed line). Data are presented as mean ± SEM. Statistical analyses were performed using unpaired t-tests or one-way ANOVA for comparisons among multiple groups. \**vs*. 6-month-old *Cort*^+/+^ or *Cort*^−/−^ MCAO saline-treated mice; ^#^*vs.* 3-month-old *Cort*^+/+^ or *Cort*^−/−^ MCAO saline-treated mice; ^&^*vs.* 6-month-old *Cort*^+/+^ MCAO saline-treated mice. *^/#/&^*p* ≤ 0.05, **^/##/&&^*p* ≤ 0.01, ***^/###/&&&^*p* ≤ 0.001, ****^/####/&&&&^*p* ≤ 0.0001.

Glial analysis revealed increased Iba1^+^ cell density and a hyper-ramified phenotype in the ischemic hemisphere of 6-month-old *Cort*^−/−^ mice, with exacerbation of age-dependent features relative to 3-month-old mice (Fig. 7C; Fig. S12A-D). Cortistatin administration restored a homeostatic microglial morphology in both genotypes. Acute astrocytic responses were also impaired in 6-month-old *Cort*^−/−^ mice, marked by disrupted glial scars and atypical astrocyte morphology (fewer junctions, reduced branching, and diminished territorial complexity), which were normalized following cortistatin administration (Fig. 7D; Fig. S12E-H).

Assessment of BBB integrity revealed no age-related differences in IgG extravasation within genotypes; however, *Cor*t^−/−^ mice exhibited significantly greater leakage than wild-type counterparts (Fig. 7E). This disruption was accompanied by reduced vascular density and vessel diameter in 6-month-old *Cort*^−/−^ mice compared to wild-type (Fig. 7F). Cortistatin administration ameliorated BBB leakage and regulated vascular remodelling in both genotypes, consistent with findings in younger mice (Figs. 7E,F; Fig. 5D). ZO-1 disruption was also reduced after cortistatin treatment (Fig. 6G). Increased BBB permeability in cortistatin-deficient mice correlated with greater immune cell infiltration, which was notably more severe than in 3-month-old counterparts. This exacerbation was fully attenuated following cortistatin treatment in both phenotypes (Fig. 6H).

Notably, the systemic immune response to stroke was amplified in this 6-month-old cohort, showing elevated IL-12, IL-6, and MCP-1, particularly in the absence of cortistatin (Fig. S13). Cortistatin treatment significantly attenuated these inflammatory mediators across genotypes.

In summary, the premature aging phenotype driven by cortistatin deficiency exacerbates neuroinflammatory responses, glial and vascular dysfunction, and peripheral immune activation following ischemia. Exogenous cortistatin confers robust neurovascular protection and immune regulation in both 3 and 6-month-old mice, underscoring its therapeutic potential across distinct developmental stages.

## DISCUSSION

Stroke is a systemic disorder characterized by intertwined neurovascular and immune mechanisms. These interactions evolve across spatiotemporal scales, shaping both tissue injury and recovery [4]. Despite progress, preclinical studies often rely on oversimplified models that insufficiently capture aging, comorbidities, and long-term outcomes, contributing to the persistent translational gap in stroke management. Defining endogenous mediators that coordinate these multi-system interactions across disease phases is therefore critical for improving both mechanistic understanding and therapeutic development.

Using cortistatin-deficient mice, we demonstrated for the first time that endogenous cortistatin limits ischemic damage and promotes recovery. Considering the heightened susceptibility of these animals to other experimental models of neural injury [14,16], we selected a mild MCAO paradigm to minimize mortality and filament-induced endothelial injury [58]. This approach yields consistent infarct sizes while enabling the detection of genotype-dependent effects. Acute (48 h) and early subacute (7 days post-stroke) phases were chosen to capture the transition from injury to repair, a critical window for determining long-term outcome and assessing therapeutic efficacy [30,31]. A major strength of this study is the incorporation of age as a biological variable. While aging is the primary risk factor for stroke vulnerability and recovery [59], most preclinical studies rely on young animals. To address this gap, we examined 3-month-old and 6-month-old mice. Our selection of the 6-month time [jc1]point, rather than the conventional 12–18-month range used in aging studies, was specifically driven by the unique biology of the cortistatin-deficient background. Preliminary data from our group described that 6-month-old mice already exhibit heightened susceptibility to neural injury and an aging-like phenotype [56], justifying the use of these specific age cohort. Furthermore, this group represents a clinically relevant population, given the rising incidence of stroke in individuals under 70 years of age [60]. While the inclusion of older mice would have provided additional insights, they were excluded from this study due to excessive mortality and reduced model reliability, which hindered their inclusion in long-term assessments.

### Cortistatin as an integrative mediator in ischemia

Here, we identify cortistatin as a conserved and critical neuroimmune modulator of stroke pathophysiology. Integrative transcriptomic analysis of human and murine datasets consistently revealed a robust downregulation of cortistatin following stroke, across tissues, models, and disease stages. This finding aligns with recent reports identifying cortistatin as a DEG in the ischemic brain [61,62], and supports its conserved involvement in cerebral ischemia. The inverse association between cortistatin expression and stroke severity suggests a causal contribution to disease progression rather than a passive consequence of injury, raising the possibility that timely restoration of cortistatin could confer neuroprotection.

Interestingly, comparative analyses in human stroke datasets confirmed that the sustained reduction in cortistatin expression in ischemic brain regions is distinct from related neuropeptides such as somatostatin and ghrelin [62,63]. Although somatostatin and neuropeptide Y-expressing interneurons are selectively preserved after ischemia [64,65], these are not substitutes of cortistatin as it exerts unique immunomodulatory and vasoregulatory roles not shared by somatostatin [6,66]. Reduced cortistatin expression has also been implicated in other neurodegenerative/neurovascular disorders, including Alzheimer’s disease [67] and diabetic retinopathy [68]. Moreover, the deletion of the chromosomal region containing the cortistatin gene in humans causes severe neurological deficits [69].

Importantly, unlike the somatostatin promoter, a preliminary *in silico* analysis of the *Cort* gene regulatory region reveals functional binding sites for NF-κB, CRE, and Hif-1α, allowing cortistatin expression to be tightly coupled to the metabolic and inflammatory state of the cell. During acute cerebral ischemia, multiple convergent mechanisms may drive the selective depletion of cortistatin. First, energy failure and HIF-1α induction in GABAergic interneurons could redirect transcription from neuropeptide biosynthesis toward hypoxic survival programs. Additionally, global cAMP depression suppresses CREB-mediated *Cort* promoter activity. Finally, the sustained cytokine storm may reprogram both neuronal and immune cells toward states incompatible with cortistatin synthesis. In fact, cortistatin secretion is significantly reduced in brain endothelial cells under oxygen-glucose deprivation-reperfusion conditions [6].

The functional impact of this depletion is magnified by the neuropeptide’s notably complex receptor pharmacology. Cortistatin binds with high affinity to all five somatostatin receptor subtypes (SSTR1–5) and the ghrelin receptor (GHSR), enabling pleiotropic actions across the neurovascular unit. Our receptor expression profiling reveals that SSTR4 and SSTR2 are the predominant cortistatin-binding receptors expressed in the post-ischemic brain, consistent with their established roles as key mediators of cortistatin’s anti-inflammatory and neuroprotective actions. At the BBB, cortistatin-driven endothelial protection is known to be mediated primarily through SSTR2 and SSTR4, which, unlike somatostatin signalling, can stimulate cAMP accumulation in endothelial cells to promote tight junction integrity, cytoskeletal stabilization, and the suppression of endothelial inflammatory activation [6]. Synergistic co-activation of SSTR5 and GHSR may further amplify these vascular protective effects through intracellular signal convergence. In parallel, GHSR-mediated signalling suppresses TLR4 and NLRP3 inflammatory cascades in microglia and endothelial cells, contributing to the preservation of neurovascular homeostasis [70]. In peripheral and infiltrating immune cells, cortistatin predominantly engages SSTR2 (constitutively expressed in monocytes and macrophages) and SSTR4 (upregulated under inflammatory conditions) to suppress pro-inflammatory cytokine production and promote resolution [9,71,72]. Although the mechanistic dissection of receptor-specific signalling pathways represents a critical avenue for future investigation, our evidence from transcriptomic profiling, i*n vitro* cell culture experiments, and receptor expression analyses supports a model in which cortistatin exerts its protective effects through Gi-coupled somatostatin receptors restraining PI3K/Akt and Ras/MAPK inflammatory cascades. In this model cortistatin may coordinate vascular and neuroimmune protection through overlapping but distinguishable receptor-effector axes, with SSTR2 and SSTR4 serving as the primary transducers of its pleiotropic protective effects in the ischemic brain. However, the potential for heterodimeric interactions among cortistatin receptors and as-yet uncharacterized receptors represent important mechanistic questions that will require targeted receptor pharmacology and conditional genetic approaches to resolve.

### Cortistatin influences ischemia outcomes

Cortistatin deficiency markedly exacerbated ischemia-induced molecular and cellular responses. Although both wild-type and cortistatin-deficient mice activated inflammatory and remodelling pathways, these responses were amplified and prolonged in the absence of cortistatin. Transcriptomic signatures revealed heightened neuroinflammation, neurovascular dysfunction, and impaired synaptic, axonal, and plasticity-related programs, consistent with maladaptive post-ischemic responses that drive secondary injury and impede regeneration [73]. Consistent with these molecular findings, cortistatin-deficient mice exhibited poorer neurological outcomes at acute stages across ages. Increased lesion size was observed in 3-month-old cortistatin-deficient mice, supporting a neuroprotective role for cortistatin. In contrast, lesion size did not further increase with age in cortistatin-deficient animals, consistent with evidence that infarct volume does not fully predict neurological outcome, particularly in the context of aging [25,74,75]. Neuronal degeneration in aged brains may converge on similar endpoints regardless of initial infarct volumes, resulting in disproportionate functional decline [76,77]. These results align with previous studies demonstrating that aging primarily impairs recovery rather than increasing lesion size [76,78,79], further confirming the effect of cortistatin deficiency on reparative pathways.

Behavioural impairments were more prevalent in cortistatin-deficient mice, although motor deficits were variable. This likely reflects the heterogeneity of post-stroke phenotypes and the limitations of traditional behavioural assays, which may lack the sensitivity to detect subtle functional changes in mild injury models [25]. Additionally, the baseline anxiety-like behaviour previously described in the absence of cortistatin, driven by chronic glucocorticoid elevation [14], may confound motor assessments and obscure genotype-dependent deficits, underscoring the need for multidimensional functional evaluation.

### Glial dysregulation, systemic inflammation and neuroendocrine-immune responses

Within the CNS, cortistatin critically constrained neuroinflammation. Microglia, the CNS primary immune responders, rapidly accumulate in the peri-infarct and infarct regions after ischemia [29]. 3-month-old cortistatin-deficient mice exhibited a hyper-reactive microglial phenotype with reduced anti-inflammatory markers (*i.e.,* CD163 and IL-10), indicating amplified neuroinflammatory response [80,81], while older cortistatin-deficient mice showed marked microglial accumulation in the infarct core and hyper-ramified morphologies likely associated with excitotoxicity [53,82] and synaptic loss [83]. Transcriptomic analysis reveals enrichment of pathways associated with microglial activation, neuroinflammatory responses, apoptosis, and axonal injury in *Cort^−/−^* mice. This is accompanied by upregulation of key reactive glial regulators (*e.g., Osm, Spp1, Fcgr1, Isg15*), enhanced phagocytic signatures (*e.g., Msr1, Cd74, Ifi30*), or activation of central microglial pathways (*e.g., Trem2-Tyrobp, Csf1r, Cx3cr1, C1qa*) supporting sustained neuroinflammation. These findings align with previous evidence of exaggerated inflammatory responses in cortistatin-deficient microglia and suggest that cortistatin limits maladaptive microglial activation, particularly with aging [56]. Interestingly, as chronic stress induces hyper-ramified microglia linked to enhanced neuronal activation and impaired spatial memory [84], the stress-like behaviour of cortistatin-deficient mice may reflect a pre-existing neuroinflammatory state that exacerbates ischemic pathology.

Astrocytes, which play central roles in neuroinflammation, BBB support, vascular dynamics, and tissue repair [85], were also profoundly dysregulated by cortistatin deficiency. In cortistatin-deficient mice, particularly in 6-month-old animals, astrocyte reactivity was biphasic, characterized by attenuated scar formation at acute stages followed by exaggerated scarring during subacute recovery. Both extreme contexts compromise neurorepair: insufficient early containment of necrotic core may allow lesion expansion, while excessive glial fibrosis obstructs revascularization and axonal regeneration [86–89]. These findings highlight a stage-dependent dysregulation of astrocytic responses and identify cortistatin as a temporal modulator of astrocytic transitions from protective to restorative phenotypes. Moreover, astrocytes lacking cortistatin also displayed atypical morphologies, characterized by small, elongated, poorly ramified forms, suggesting structural instability and disrupted maturation or polarization. These features were present at both ages, indicating that cortistatin loss may accelerate age-related astrocytic dysfunction. Recent studies demonstrating induction of pro-inflammatory astrocyte states in aged ischemic brains further support this hypothesis [61]. These morphological alterations correlated with disturbed glial transcriptomic landscapes in cortistatin-deficient animals, defined by the simultaneous presence of activated/deactivated microglial signatures and a phenotypic switch in astrocytes from homeostatic-protective to reactive-neurotoxic states. Such changes are predicted to impair metabolic support, glutamate clearance, and BBB integrity [73], thereby exacerbating post-ischemic dysfunction. Transcriptomic evidence of reduced oligodendrocyte maturation and myelination further indicates that cortistatin promotes white matter repair, essential for cognitive and motor recovery [90].

It is well described that stroke induces not only focal brain injury but also triggers a systemic immunocompromised state, characterized by lymphoid organ atrophy and infection susceptibility [91]. In this context, cortistatin deficiency intensified inflammatory transcriptional programs. Cortistatin-deficient mice displayed markedly increased immune infiltration into the ischemic brain, as evidenced by histological analysis and transcriptomic deconvolution, *i.e.,* enrichment of innate immune receptors (*e.g., Tlr* family members), inflammatory mediators (*Tnf, Il33*), and chemokines (*Ccl7, Ccl9, Ccl12*). These results mirror previous evidence of enhanced leukocyte transmigration across cortistatin-deficient endothelium [6], and position cortistatin as a molecular brake on leukocyte trafficking and vascular inflammation. Although the precise mechanisms remain undefined, cortistatin may regulate immune trafficking through effects on endothelial adhesion molecules, as cortistatin-deficient brain endothelial cells upregulate integrins (*Itga3*, *Itga4*) implicated in leukocyte adhesion and migration [6]. Cortistatin-deficient mice also showed heightened pro-inflammatory cytokine production, characterized by elevated IL-6, IL-12, and MCP-1, alongside reduced anti-inflammatory signalling. These factors are strongly associated with infarct expansion, neurological deterioration, and poor long-term outcome in stroke patients [92–96], reinforcing the clinical relevance of our findings. Finally, reduced spleen size and cellularity indicate systemic immune dysregulation linked to poor prognosis [97].

Additionally, at the systemic level, endocrine dysregulation likely contributes to cortistatin deficiency phenotypes. The hypothalamic–pituitary–adrenal (HPA) axis is a well-known coordinator of stress and immune responses, with studies establishing that elevated glucocorticoid levels correlate with increased mortality and worse recovery in both humans and experimental models [98–101]. It has also been reported that cortistatin absence chronically elevates glucocorticoids [12], which likely increases stroke susceptibility. Notably, clinical data indicate that cortistatin treatment in humans attenuates HPA axis hyperactivation, suggesting a conserved regulatory role [102]. Despite this chronic glucocorticoid elevation, cortistatin-deficient glia and endothelial cells exhibited pro-inflammatory phenotypes resistant to glucocorticoid-mediated suppression. This paradox may reflect altered downstream signalling, supported by reduced expression of the glucocorticoid-responsive endothelial factor *Anxa1*, which is implicated in BBB regulation [6,103]. Thus, cortistatin emerges as a critical modulator of neuroendocrine-immune crosstalk.

### BBB disruption, endothelial dysfunction, and vascular aging phenotype

At the vascular level, cortistatin preserves tight junction integrity, whereas its deficiency promotes vascular dysfunction and extracellular matrix remodelling (*e.g., Adamts1, Thbs1*) and stress-response pathway activation (*e.g., Hmox1, Nfe2l2, Stat3*). BBB disruption was indeed markedly exacerbated in cortistatin-deficient mice, as evidenced by increased IgG extravasation. This was evident in both 3-month and 6-month-old deficient animals following acute ischemia, extending previous findings of heightened BBB vulnerability under inflammatory conditions [6]. Brain endothelial cells lacking cortistatin displayed intrinsic permeability defects and amplified sensitivity to ischemic-like conditions. Consistent with this, reduced cortistatin expression in mouse and human endothelial cells following oxygen-glucose deprivation supports its direct role in endothelial resilience [6]. Notably, 6-month-old cortistatin-deficient mice showed reduced vascular density relative to young counterparts, a difference absent in wild-type mice, indicating that cortistatin deficiency accelerates premature cerebrovascular decline. Indeed, younger deficient mice recapitulated many molecular and pathological features previously described for older wild-type animals (*e.g.,* increased inflammation, interferon signalling, disrupted synaptic maintenance, exaggerated glial activation, and BBB breakdown) [61]. These data demonstrate a synergistic interaction between cortistatin loss and accelerated aging in driving neurovascular and neuroimmune decline.

### Therapeutic restoration of cortistatin and translational relevance

Considering the narrow therapeutic window of current reperfusion strategies, identifying effective interventions during the early post-stroke period is critical, as it can determine long-term outcome [2]. Our data demonstrate that cortistatin deficiency disrupts the early activation of pathways governing inflammation resolution, tissue repair, neurogenesis, synaptic plasticity, and cognitive function, resulting in defective recovery by 7 days post-injury. Specifically, this manifests as reduced neuronal repopulation, compromised axonal integrity and myelination, aberrant glial scarring, and reduced VEGF expression, all of which severely impair neurovascular remodelling and repair [104]. Interestingly, studies from our group show that cortistatin-deficient mice exhibit myelin abnormalities in multiple sclerosis models [105], supporting a broader role for cortistatin in axonal integrity and neurorepair, though mechanisms remain unclear.

Critically, restoration of cortistatin levels conferred robust neuroprotection across multiple dimensions. Acute administration reduced brain injury and neurological deficits, normalized glial and immune responses, preserved BBB integrity, and enhanced neurotrophic factor expression, including BDNF and GDNF, which are known mediators of neurogenesis and functional recovery [106,107]. These protective effects are consistent with the broad expression of cortistatin across neurovascular cell types (including GABAergic interneurons, microglia, astrocytes, pericytes, and endothelial cells) as well as across peripheral immune compartments, underscoring its pleiotropic nature. Cortistatin exerts protection through two complementary and temporally coordinated mechanisms that combines direct actions on resident neurovascular cells and indirect effects derived from systemic immunomodulatory regulation [72], which limits peripheral immune infiltration and promotes inflammatory resolution. Rather than operating independently, these local and systemic arms act synergistically, reinforcing one another across the stroke continuum. While overall functional outcomes were improved after cortistatin treatment, the effect on motor recovery was limited in 6-month-old cortistatin-deficient mice. This may reflect age-related dopaminergic dysregulation or reduced motor drive inherent to this genotype. Given the known influence of cortistatin on motivational and locomotor circuits [10], further studies should dissect its role in corticostriatal circuitry.

Importantly, while some studies indicate that broad immune suppression may increase infection susceptibility, complicating the risk-benefit profile of anti-inflammatory therapies [108], our findings demonstrate that cortistatin exerts a context-dependent, rather than indiscriminate, modulation of glial and immune responses. This aligns with the emerging consensus that effective therapeutic strategies must be temporally and contextually tailored [19,109]. Notably, these therapeutic benefits extended into the subacute phase and remained consistent across age groups, highlighting a mechanism capable of overcoming age-related resistance to neuroimmune therapies.

Taken together, these findings establish cortistatin as a non-redundant homeostatic factor at the interface of the neuroimmune axis, whose deficiency initiates a self-amplifying inflammatory cascade that progressively undermines neurovascular integrity, neuronal survival, and functional recovery. Further exploration of cortistatin analogues or specific receptor agonists may yield innovative therapies aimed at restoring neuroimmune equilibrium and promoting a sustained recovery after ischemic injury.

## CONCLUSIONS

Collectively, this study advances our understanding of stroke pathophysiology by repositioning stroke as a multisystem disorder rather than a localized cerebrovascular insult. Our results establish for the first time cortistatin as a pleiotropic endogenous regulator in stroke, associated with the modulation of immune responses, glial homeostasis, BBB stability, and regenerative signalling across the injury-repair continuum. Its deficiency triggers uncontrolled inflammation and vascular failure, recapitulating an aging-like neurovascular decline, and impairing recovery. Conversely, cortistatin supplementation confers neuroprotection and promotes repair across different disease stages, extending the therapeutic window beyond current time-limited interventions. The translational relevance of these findings is further reinforced by the consistent downregulation of cortistatin in human stroke patients, positioning it as both a potential biomarker and a promising avenue for therapeutic intervention after ischemic injury.

## STUDY LIMITATIONS AND FURTHER DIRECTIONS

Certain limitations should be acknowledged in the context of future research, without diminishing the robustness or significance of the present findings. First, experiments were conducted exclusively in male mice; incorporating female cohorts in subsequent studies will be essential to account for known sex-dependent differences in stroke recovery and validate the generalizability of these results. Additionally, the intrinsic anxiety phenotype of cortistatin-deficient mice may introduce confounds in behavioural tasks, and observed functional trends in our cohorts should be interpreted with caution. Second, the 6-month-old knockout mice do not represent an advanced ageing model *per se*; rather, this timepoint was strategically selected to investigate the early-aging phenotype inherent to cortistatin deficiency, and reflects the upper feasible limit for stroke induction in this genotype given the prohibitively high post-MCAO mortality observed at 8–12 months. Future studies using inducible or cell-specific conditional models may help circumvent this constraint. Third, while our study successfully identifies the pleiotropic effects of cortistatin and several key molecular mediators, the precise downstream signalling cascades activated upon receptor engagement in different brain and immune cell populations remain to be fully dissected. Finally, although bulk RNA-seq combined with orthogonal immunohistochemistry provides a comprehensive transcriptomic and structural landscape, future studies employing single-cell RNA sequencing will be invaluable to resolve cell-type-specific responses and more precisely identify the primary cellular targets of cortistatin.

## ABBREVIATIONS

BBB: blood-brain barrier
BECs: brain endothelial cells
BP: biological processes
BSA: bovine serum albumin
CC: cellular components
CD31/PECAM-1: cluster of differentiation 31
CD45: cluster of differentiation 45
CL: contralateral
CNS: central nervous system
CST: cortistatin
CV: cresyl violet
CVST: cerebral venous sinus thrombosis
DEGs: differential-expressed genes
ECs: endothelial cells
ECM: extracellular matrix
FDR: false discovery rate
GEO: gene expression omnibus database
GFAP: glial fibrillary acidic protein
GHSR: ghrelin receptor
GO: gene ontology
GSEA: gene set enrichment analysis
HPA: hypothalamic–pituitary–adrenal
Iba1: ionized calcium-binding adapter molecule 1
log2FC: log2 fold change
MAP2: microtubule-associated protein 2
MBP: myelin basic protein
MCA: middle cerebral artery
MCAO: middle cerebral artery occlusion
MF: molecular functions
MGPs: marker gene profiles
NPC: neural progenitor cells
PBMCs: peripheral blood mononuclear cells
PCoA: principal coordinates analysis
PT: photothrombotic
rCBF: regional cerebral blood flow
rtPA: recombinant tissue plasminogen activator
RPKM: reads per kilobase of exon per million mapped fragments
STAIR: stroke therapy academic industry roundtable
TJ: tight-junctions
TNF-α: tumour necrosis factor-alpha
TPM: transcript-per-million
VEGF: vascular endothelial growth factor
ZO-1: zonula occludens-1

## ACKNOWLEDGEMENTS

The authors would like to thank the Animal Care Unit and Microscopy Service Unit staff from the IPBLN for their technical support. We also acknowledge support from the PTI+ NEUROAGING, Consejo Superior de Investigaciones Científicas (CSIC), Spain.

## AUTHOR CONTRIBUTIONS

E.G-R conceived, designed, and supervised the entire study. J.C-G and E.G-R outlined the experiments and analysed all the data. J.C-G, P.V-R, I.S-M, A.C-M, and I.F-L performed the experiments. P.V-R and JL. R analysed and interpreted the transcriptomic data. J.C-G, L.B, M.P, L.H, and E.G-R contributed to data interpretation and discussion of the results. J.C-G and E.G-R wrote and revised the manuscript. All authors have read and approved the final version of the manuscript.

## FUNDING

This work was supported by the Spanish Ministry of Economy and Competitiveness (MCIN)/Spanish Research Agency (AEI)/10.13039/501100011033 grants: SAF2017-85602-R and PID2020-119638RB-I00 (to E.G-R.), FPU-program FPU17/02616 and Ramón Areces Foundation postdoctoral fellowship (to J.C-G.), FPI-program PRE2018-084824 (to I.S-M.), PRE2021-100172 (to P.V-R.) and FPU-program FPU23/00152 (to A.C-M); the grant PROYEXCEL_00867 from the Regional Andalusian Government (to E.G.-R.); the Swiss Science Foundation FNS 310030_212233 grant to L.H

## DATA AVAILABILITY

RNA-seq data are available in the GEO repository under accession number GSE302342, https://www.ncbi.nlm.nih.gov/geo/query/acc.cgi?acc=GSE302342.

## DECLARATIONS

### Ethics approval and consent to participate

*Institutional Animal Care and Use Committee (IACUC):* All experimental protocols employed in this study were approved by the Animal Care and Use Board and the Ethical Committee of the Spanish National Research Council (Animal Care Unit Committee IPBLN-CSIC # protocol CEEA OCT/2017.EGR) and conducted in accordance with Directive 2010/63/EU of the European Parliament on the protection of animals used for scientific purposes.

This study involves the re-analysis of publicly available human data. The original studies from which the data were obtained received ethical approval from their respective Institutional Review Boards. Therefore, a new ethical approval or informed consent was not required for this manuscript, in accordance with local legislation and institutional requirements regarding the secondary use of anonymous public data.

### Consent for publication

Not applicable

### Competing interests

The authors declare that they have no competing interests.

### Author details

^1^Institute of Parasitology and Biomedicine (IPBLN-CSIC), Granada, Spain; ^2^University of Lausanne, Lausanne, Switzerland; ^3^Lausanne University Hospital, Lausanne, Switzerland; ^4^Functional Genomics Center Zurich, ETH Zurich and University of Zurich, Zurich, Switzerland; ^5^Swiss Institute of Bioinformatics, Amphipôle, Quartier UNIL-Sorge, Lausanne, Switzerland

## Notes

### Competing Interest Statement

The authors have declared no competing interest.

### Summary of Updates

This version has been updated with new data, corrected typos from the previous version, improved explanations of certain aspects, and a discussion of the limitations of the study.

