## Additional File 2 for "Lack of cortistatin drives neuroimmune and vascular dysfunction in brain ischemia"

### A. Extended Methods

#### 1. Next-generation transcriptome sequencing (RNA-seq) and data analysis.

The *reanalyzerGSE* pipeline v.3.0.0 (Andrés-León and Rojas, 2019), which builds upon *miARma-Seq* (Nikolayeva and Robinson, 2014), was used to conduct quality control, read alignment, and quantification. To identify differentially expressed genes (DEGs), we employed the *edgeR* R package (Reeb et al., 2015). Analyses were carried out using default parameters, except for a customized threshold applied during the filtering of lowly expressed genes such as cortistatin. We filtered out genes with zero read counts in all but one sample. The reference genome and annotation used were *Mus musculus* GRCm39 primary assembly and GENCODE annotation version M28, respectively. The average proportion of uniquely mapped reads across samples was 78.5%. Gene expression levels were normalized and quantified using both RPKM (Reads Per Kilobase Million) and TPM (transcript-per-million). RPKM was used for all the analyses, except for the marker gene profile (MGP) studies, which were based on TPM, as detailed below. To evaluate data replicability and assess the biological variability, we applied Principal Coordinates Analysis (PCoA) and unsupervised hierarchical clustering on normalized counts (Ritchie et al., 2015; Brionne et al., 2019). In comparisons where a large variability could potentially mask differential expression by separating biological replicates into distinct groups, only consistent and clearly grouped replicates were included. Nevertheless, all analyses retained three or more biological replicates. For functional enrichment analyses, we used the *enrichGO* function of the ClusterProfiler package v4.12.6 (Yu et al., 2012) with default parameters, except for using as background the whole universe of transcripts with quantified expression. We identified significantly enriched Gene Ontology (GO) terms (adjusted p-value < 0.05) related to Biological Processes (BP), Molecular Functions (MF), and Cellular Components (CC) for each DEGs group. Additionally, for the limited DEGs identified in the comparisons *Cort*$^{-/-}$ *vs Cort*$^{+/+}$, we applied a Gene Set Enrichment Analysis (GSEA) using the package fgsea v1.30.0. Genes were ranked according to their log2 fold change (log2FC), default parameters were used, except for 1,000 permutations. Only gene sets containing between 5 and 500 genes were included. Gene sets with an adjusted p-value < 0.05 were considered significantly enriched and reported. GSEA was run for GO, KEGG, Reactome, and Hallmark databases from MsigDB (Castanza et al., 2023; Subramanian et al., 2005). Unless indicated otherwise, all statistical analyses and visualizations were performed in R v4.4.1 using base functions (Huber et al., 2015), Bioconductor packages (Wickham, 2011), and ggplot2 v3.5.1 (Schep and Kummerfeld, 2017; Wickham, 2016) along with its extensions.

#### 2. Cell-specific gene sets and cell type proportion estimation

To analyse cell type proportions in the experimental groups from our study, we performed a computational deconvolution. For this, we used TPM normalized gene expression data as input for the markerGeneProfile R package v1.0.4 (REF 11) and estimated marker gene profiles (MGPs) for various cell types. To reduce transcriptional noise, a filtering step was first applied, and genes with expression values below the median across all samples were excluded. The estimation was performed using the mgpEstimate function, during which genes showing low correlation to the majority of other marker genes were removed, following the methodology described by Mancarci et al. (Mancarci et al., 2017). A small subset of marker gene sets was excluded due to a high proportion of removed genes (> 40 %) and/or a low proportion of variance explained by the first principal component (< 50 %). Finally, the resulting MGP values were normalized between 0 (minimum) and 1 (maximum). To assess differences in normalized MGP values between groups, a linear model (function *lm*) was fitted with normalized MGP as the dependent variable and condition (experimental groups) as the independent factor. An analysis of variance (ANOVA) was performed to evaluate the overall effect of condition on MGP values. Pairwise comparisons between levels of the condition factor were conducted using the *emmeans* R package (v1.11.0) (Androvic et al., 2020). P-values from these comparisons were adjusted using the Holm method to control for multiple testing.

**3. Integration of human and mouse datasets for cortistatin gene expression analysis**

We leveraged results from previously published studies to examine cortistatin expression at the transcriptional level. Datasets were downloaded from the Gene Expression Omnibus (GEO) database (Clough et al., 2024). Regarding human transcriptional profiles of blood cells, studies collected samples from different populations: (i) peripheral blood mononuclear cells (PBMCs) from acute (24 h post-onset), subacute (7 days post-onset) ischemic stroke, and healthy individuals (GSE122709); (ii) PBMCs from patients with large artery occlusion undergoing interventional thrombectomy, with or without subsequent intracranial haemorrhage (GSE199435); (iii) and peripheral leukocytes from patients with minor (NIHSS score < 8) and with severe (NIHSS score ≥ 8) acute ischemic stroke on admission, compared to healthy controls (GSE202709). One study included human brain tissue with the temporal lobe dissected from the post-mortem brain of acute ischemic stroke patients and matched contralateral healthy tissue as control (GSE56267). Mice datasets included different experimental stroke models, namely: (i) transient MCAO (60 min) with samples from peri-infarct cortex (GSE253799) and penumbral tissue (GSE227186) analysed together with sham-operated mice in the acute phase (24 h reperfusion and sacrifice); (ii) transient MCAO (60 min) with samples from ipsilateral cortex from sham and ischemic mice sacrificed at 24 h (acute phase) or 28 days after stroke (chronic phase) (GSE131712); cerebral venous sinus thrombosis (CVST) with samples of the cerebral cortex of sham mice and ischemic animals, analysed at 2 and 7days post-injury (GSE245455); and (iii) photothrombotic (PT) stroke models with peri-infarct cortex obtained at the acute stage (3 days post-injury) (GSE261613); with peri-infarct and infarct core regions obtained at the subacute stage (7 days post-injury) (GSE280846); and with contralateral and peri-infarct cortex at 7- and 14-days post-stroke (GSE184917). Cortistatin gene expression levels were reported as normalized RPKM, FPKM, TPM, or count values. The data from GSE122709 and GSE56267 were generated and analysed with GEO2R (Clough et al., 2024), and the rest was provided by the authors in the original publication

### B. Extended Results

A

B

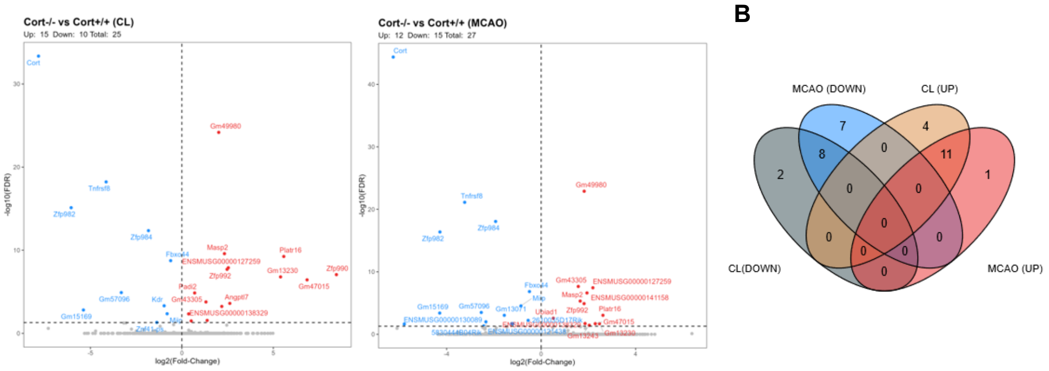


C

| KEGG Pathways | NES | p.adj |
|---|---|---|
| NOD-like receptor signaling pathway | 1.706 | 0.003 |
| Epstein-Barr virus infection | 1.606 | 0.005 |
| Herpes simplex virus 1 infection | 1.605 | 0.006 |
| Cytosolic DNA-sensing pathway | 1.668 | 0.013 |
| RIG-I-like receptor signaling pathway | 1.681 | 0.015 |
| Phagosome | 1.605 | 0.015 |
| Human papillomavirus infection | 1.428 | 0.015 |
| Hepatitis B | 1.585 | 0.018 |
| Human cytomegalovirus infection | 1.475 | 0.022 |
| Hepatitis C | 1.567 | 0.023 |

| Biological Processes GO Pathways | NES | p.adj |
|---|---|---|
| Positive regulation of innate immune response | 1.611 | 0.001 |
| Positive regulation of response to biotic stimulus | 1.594 | 0.001 |
| Regulation of innate immune response | 1.560 | 0.001 |
| Pattern recognition receptor signaling pathway | 1.676 | 0.002 |
| Activation of innate immune response | 1.635 | 0.002 |
| Cytoplasmic pattern recognition receptor signaling pathway | 1.713 | 0.006 |
| Innate immune response-activating signaling pathway | 1.622 | 0.006 |
| Negative regulation of response to biotic stimulus | 1.723 | 0.010 |

| Hallmark gene sets Pathways | NES | p.adj |
|---|---|---|
| Interferon gamma response | 1.534 | 0.032 |
| E2F Targets | 1.418 | 0.032 |
| IL6-JAK-STAT3 Signaling | 1.540 | 0.035 |
| G2M Checkpoint | 1.418 | 0.035 |
| TNFa signaling via NFkB | 1.378 | 0.043 |
| KRAS signaling UP | 1.345 | 0.043 |

| Reactome Pathways | NES | p.adj |
|---|---|---|
| NABA ECM Regulators | 1.593 | 0.034 |

**Figure S1. Characterization of molecular responses in the ischemic brain of cortistatin-deficient mice.** Differential gene expression analysis was conducted in *Cort*$^{-/-}$ *vs Cort*$^{+/+}$ mouse brains 48 h after ischemic stroke. **(A)** Volcano plots display differentially expressed genes (DEGs) between *Cort*$^{-/-}$ *vs Cort*$^{+/+}$ in both uninjured contralateral (CL, left) and ischemic (right) hemispheres following 20 min MCAO. Each dot represents one gene. Significantly upregulated (red) and downregulated (blue) genes (with false discovery rate, FDR, $p < 0.05$) for cortistatin-deficient mice are shown. Non-significant genes are shown in grey. Full description of DEGs is detailed in Additional file 1: Tables S2 and S3. **(B)** Venn diagram showing the overlap of significantly upregulated and downregulated genes in the healthy and ischemic hemispheres. **(C)** Functional enrichment analysis (GSEA) was performed using multiple databases, including KEGG, Gene Ontology (GO), Reactome, and Hallmark pathways. Details in Additional file 1: Table S4. ECM: extracellular matrix; JAK-STAT: Janus kinases-signal transducer and activator of transcription; KRAS: Kirsten rat sarcoma virus; KEGG: Kyoto Encyclopedia of Genes and Genomes; NES: normalized enrichment score; NOD: nucleotide-binding oligomerization domain; RIG: retinoic acid-inducible gene I. NABA: extracellular matrix pathway gene set (see (Naba et al., 2012)).

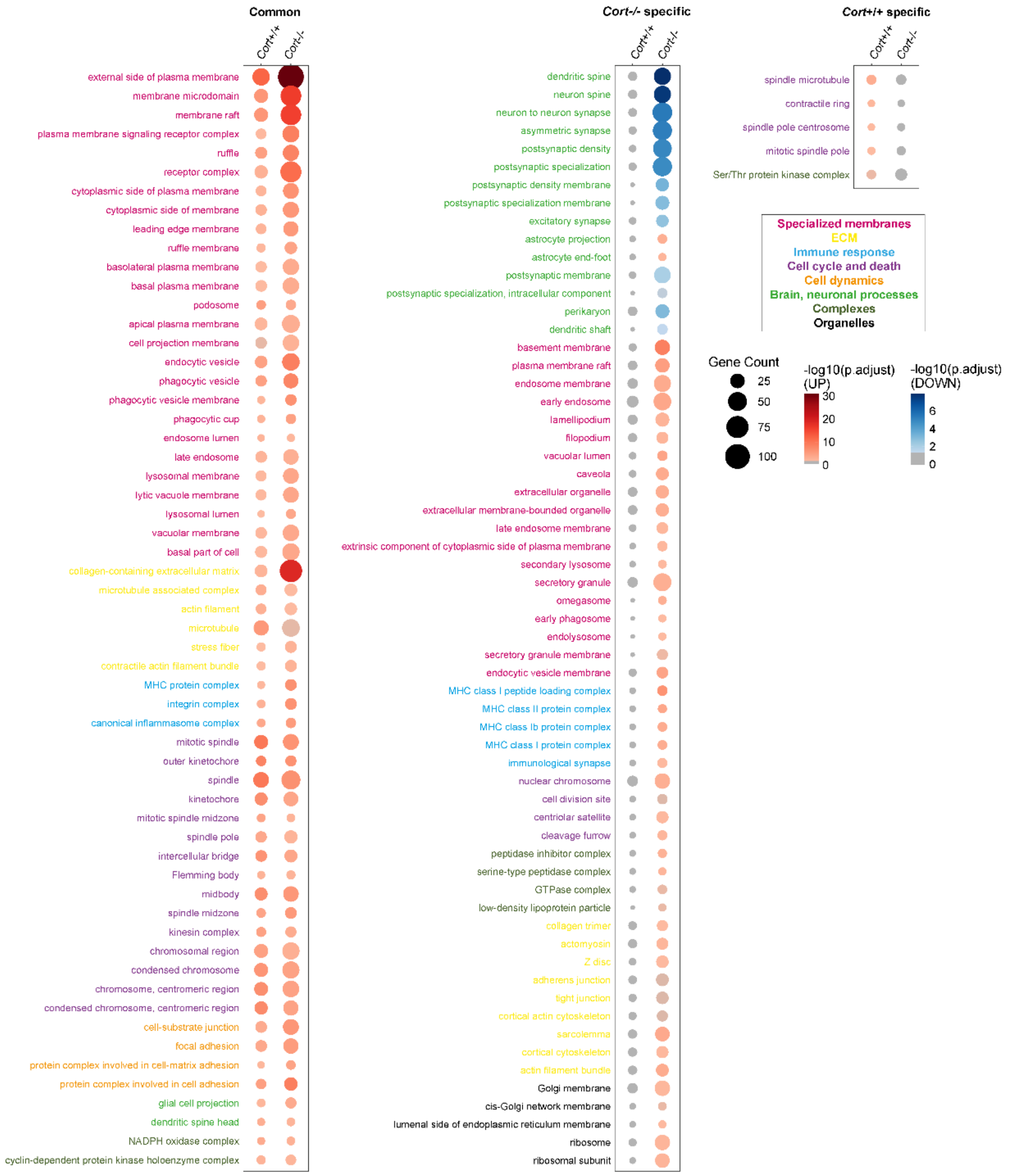


**Figure S2. Enrichment of cellular component-related pathways in cortistatin-deficient brains following ischemia.** Gene ontology (GO) enrichment analysis for cellular components was performed to identify significantly overrepresented pathways in *Cort*$^{-/-}$ compared to *Cort*$^{+/+}$ during the transition from healthy to ischemic states. Pathways were grouped into biologically related categories and manually annotated into functional networks including: specialized membranes (magenta), extracellular matrix (yellow), immune response (blue), cell cycle and death (purple), cellular dynamics (orange), brain and neuronal processes (green), complexes (olive green), and organelles (black). Shared and genotype-specific pathways are indicated. All DEGs associated with each GO term are listed in Additional file 1: Table S8. Colour intensity denotes significance (red: upregulated, blue: downregulated, grey: non-significant). Circle sizes reflect the number of DEGs in each pathway.

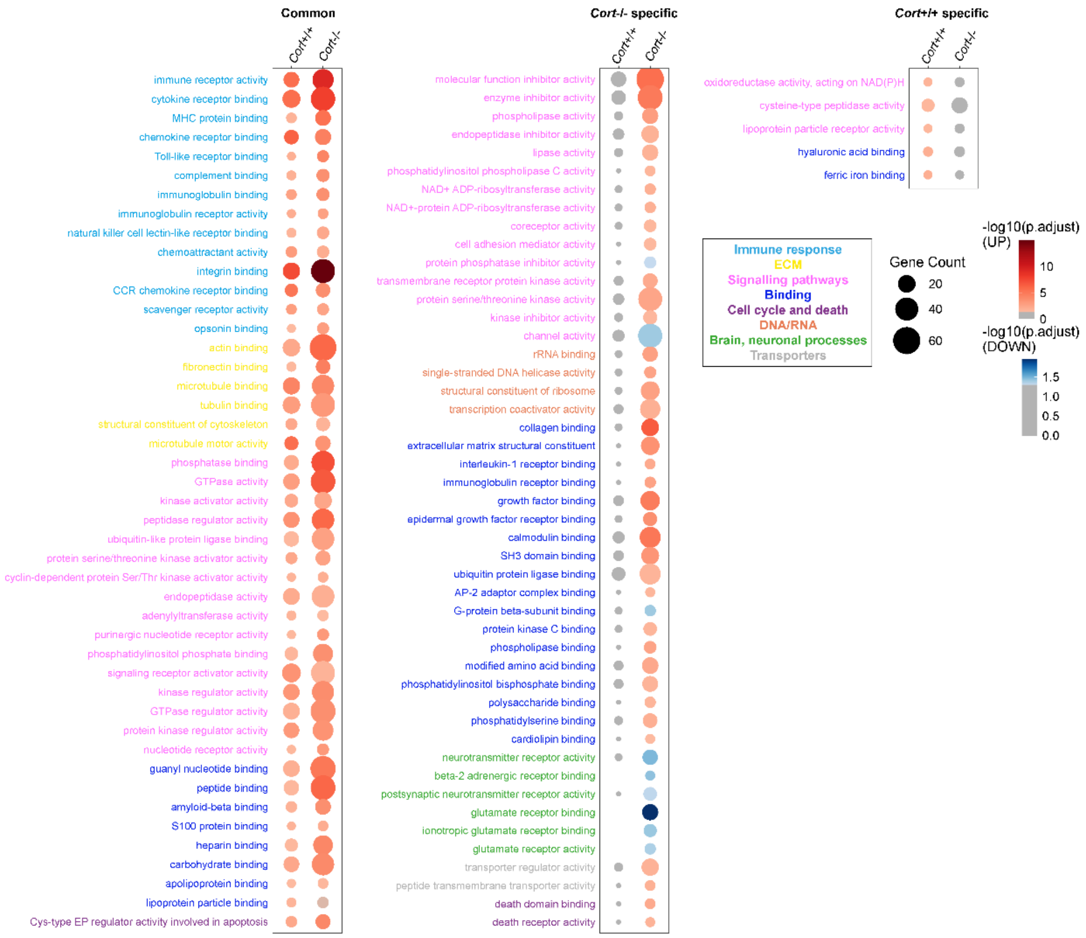


**Figure S3. Enrichment of molecular function-related ontology terms in cortistatin-deficient brains following ischemia.** Gene ontology (GO) enrichment analysis of molecular function terms was performed to identify significantly overrepresented GO molecular function terms in *Cort*$^{-/-}$ compared to *Cort*$^{+/+}$ during the transition from healthy to ischemic states. Pathways were grouped into biologically related categories and manually annotated into functional networks including: immune response (blue), extracellular matrix (ECM) (yellow), signalling (pink), molecular binding (blue), cell cycle and death (purple), DNA/RNA metabolism (orange), brain and neuronal processes (green), and transporters (grey). Shared and genotype-specific ontology terms are indicated. All DEGs associated with each GO category are listed in Additional file 1: Table S9. Colour intensity denotes significance of enrichment (red: upregulated, blue: downregulated, grey: non-significant). Circle sizes reflect the number of DEGs in each pathway.

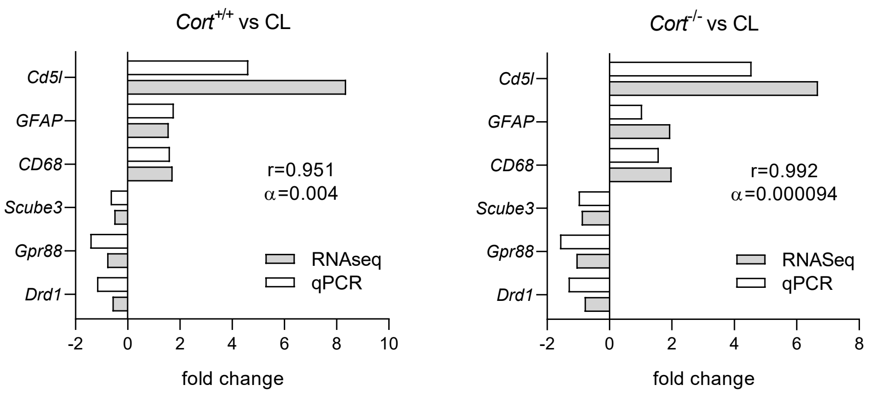


**Figure S4. Validation of RNA sequencing (RNA-seq) data by real-time qPCR**. Expression levels of selected upregulated and downregulated differentially expressed genes (DEGs) were assessed in *Cort+/+* (left) and *Cort-/-* (right) mice, comparing ipsilateral versus contralateral (CL) hemispheres 48 h after MCAO. Fold-change values obtained by RNA-seq and qPCR showed a strong and significant correlation, as determined by Spearman's rank correlation analysis. N = 3–6 samples per group.

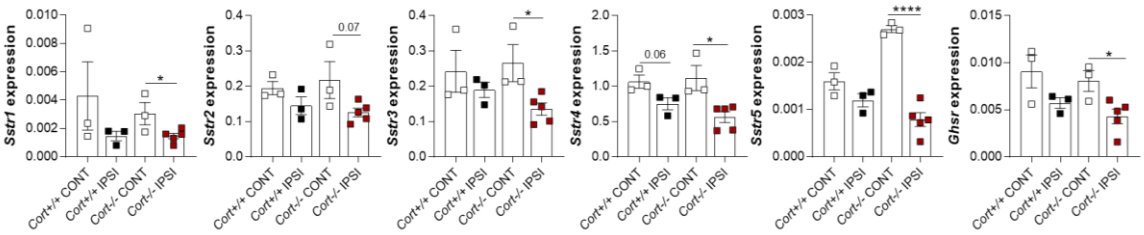


**Figure S5. Stroke provokes the downregulation of cortistatin receptors.** MCAO was performed in *Cort+/+* and *Cort−/−* mice, and animals were sacrificed at 48 h. Gene expression of cortistatin receptors (*Sstr1, Sstr2, Sstr3, Sstr4, Sstr5*, and *Ghsr*) in contralateral (CL) and ipsilateral (IP) hemispheres, assessed by qPCR and normalized to *Rplp0*. N = 3–5 mice per group. Data are presented as mean ± SEM, with dots representing individual mice. Statistical analyses were performed using unpaired t-tests or one-way ANOVA for comparisons among multiple groups. *vs. $Cort^{-/-}$ MCAO saline-treated mice. $^{*}p \le 0.05$, $^{****}p \le 0.0001$.

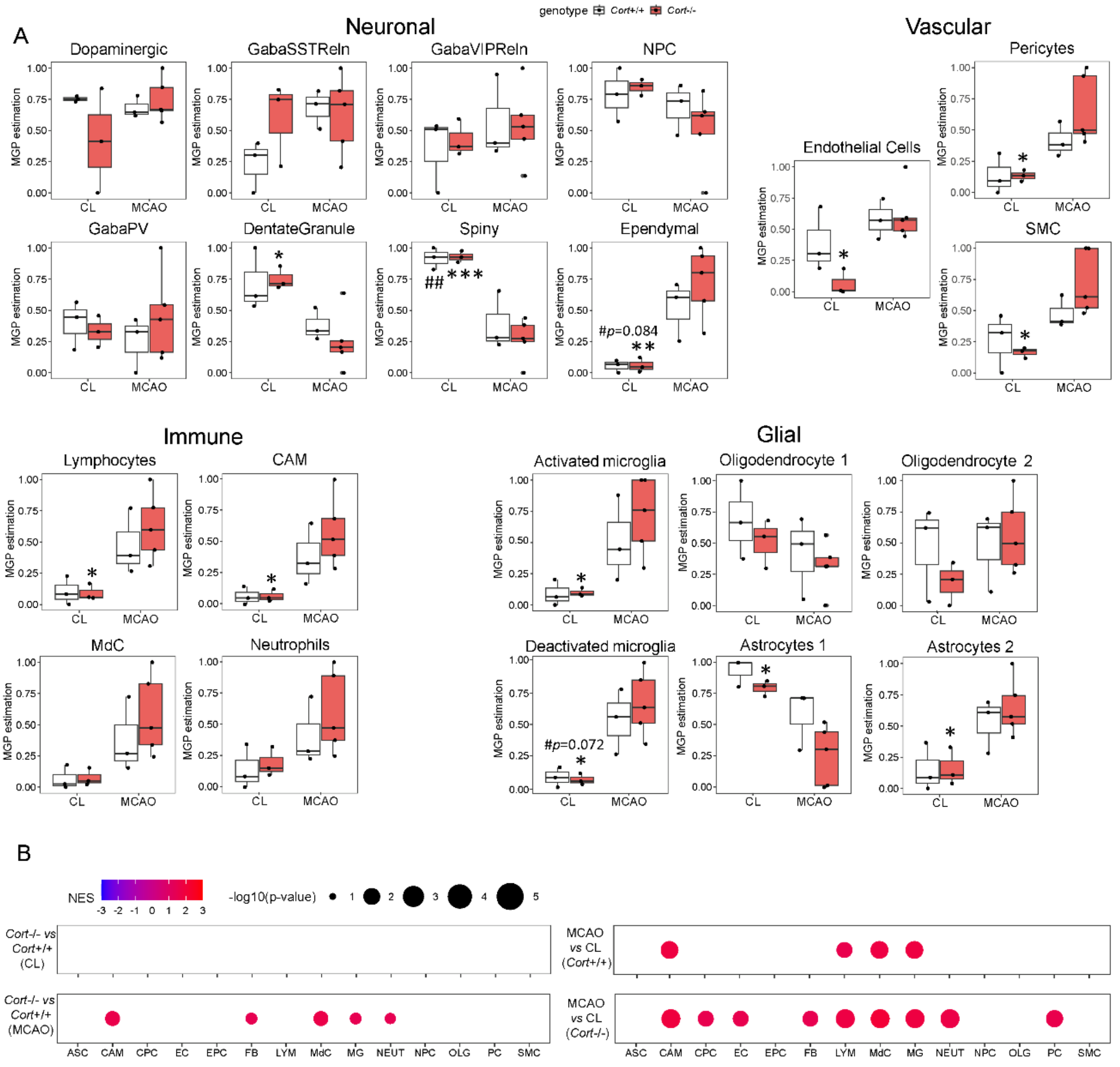


**Figure S6. Cortistatin deficiency alters brain cell marker proportions following ischemic stroke.** Cell-type-specific gene expression patterns were identified by deconvolution of bulk RNA transcriptome data as described in Methods. **(A)** Boxplots display changes in cell-type composition between healthy (CL) and ischemic (MCAO) hemispheres isolated from 12-weeks-old $Cort^{-/-}$ and $Cort^{+/+}$ animals 48 h post-stroke. Marker gene profile (MGP) analysis was used to predict cell population proportions based on known markers. Cell signatures were grouped into four main categories: neuronal (including dopaminergic, GABAergic-somatostatin (SST), GABAergic-vasoactive intestinal peptide (VIP), and parvalbumin (PV)-expressing interneurons, neural progenitor cells, and dentate granule, spiny and ependymal neurons), vascular (including endothelial cells, pericytes, and smooth muscle cells), immune (including lymphocytes, central nervous system-associated macrophages, monocyte-derived cells, and neutrophils) and glial cell types (including microglia, astrocytes, and oligodendrocytes). Statistical analyses were performed using unpaired t-tests or one-way ANOVA for comparisons among multiple groups. *$vs.$ $Cort^{-/-}$ MCAO saline-treated mice, #$vs.$ $Cort^{+/+}$ MCAO saline-treated mice, $^{*/\#}p \le 0.05$, $^{**/\#\#}p \le 0.01$, $^{***/\#\#\#}p \le 0.001$. **(B)** Comparative GSEA performed using cell-type–specific gene sets (Zheng et al., 2022) between $Cort^{-/-}$ and $Cort^{+/+}$ in healthy *vs.* ischemic hemispheres (left) and across ischemic progression within each genotype (right). Colour corresponds with Normalized Enrichment Score (NES) and circle size indicates -log10(p-value). ASC: astrocyte, CAM: central nervous system (CNS)-associated macrophage, CPC: choroid plexus capillary endothelial cell, EC: endothelial cell, EPC: ependymocyte, FB: fibroblast, LYM: lymphocyte, MdC: monocyte-derived cell, MG: microglia, NEUT: neutrophil, NPC: neural progenitor cell, OLG: oligodendrocyte, PC: pericyte, SMC: vascular smooth muscle cell.

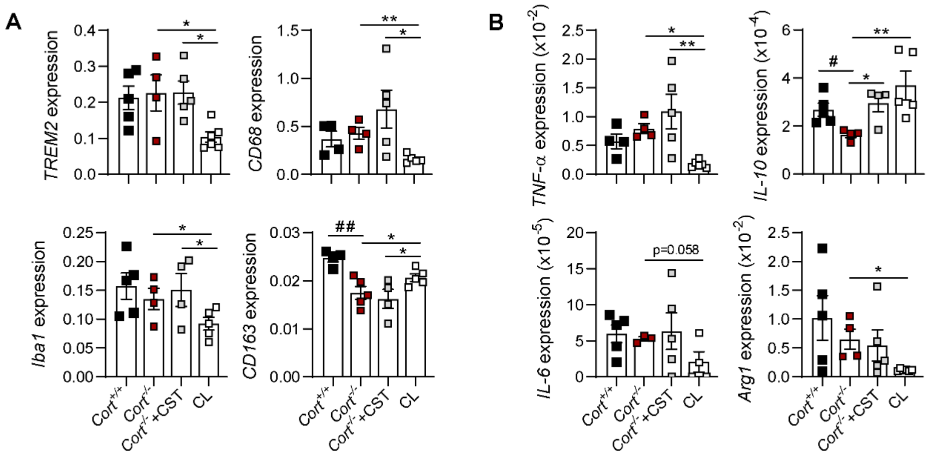


**Figure S7. Lack of cortistatin enhances post-stroke neuroinflammation.** MCAO was conducted in *Cort*$^{-/-}$ mice treated with either saline or cortistatin (144µg/kg) 24 h after reperfusion and sacrificed at 48 h. Mice were compared to *Cort*$^{+/+}$ MCAO saline-treated mice. **(A,B)** Gene expression of *Trem2, CD68, Iba1,* and *CD163* **(A)** and *TNF-α*, *IL-6*, *Arg1,* and *IL-10* **(B)** in contralateral (CL) and ipsilateral hemispheres of *Cort*$^{-/-}$ mice (MCAO saline *vs.* MCAO+CST), assessed by qPCR and normalized to *Rplp0*. N=4-6 mice/group. Data: mean ± SEM; dots represent individual mice. Statistical analyses were performed using unpaired t-tests or one-way ANOVA for comparisons among multiple groups. *$vs.$ *Cort*$^{-/-}$ MCAO saline-treated mice, $^{\#}vs.$ *Cort*$^{+/+}$ MCAO saline-treated mice. $^{*/\#}p \leq 0.05$, $^{**/\#\#}p \leq 0.01$. *Arg1*: arginase 1; *CD68*: cluster of differentiation 68; *CD163*: cluster of differentiation 163; CST: cortistatin; *Iba1*: ionized calcium-binding adaptor molecule 1; *IL-6*: interleukin 6; *IL-1β*: interleukin 1β; *IL-10*: interleukin 10; MCAO: middle cerebral artery occlusion; *MCP-1*: Monocyte Chemoattractant Protein- 1; *TNF-α*: tumour necrosis factor-α; *Trem2*: triggering receptor expressed on myeloid cells 2.

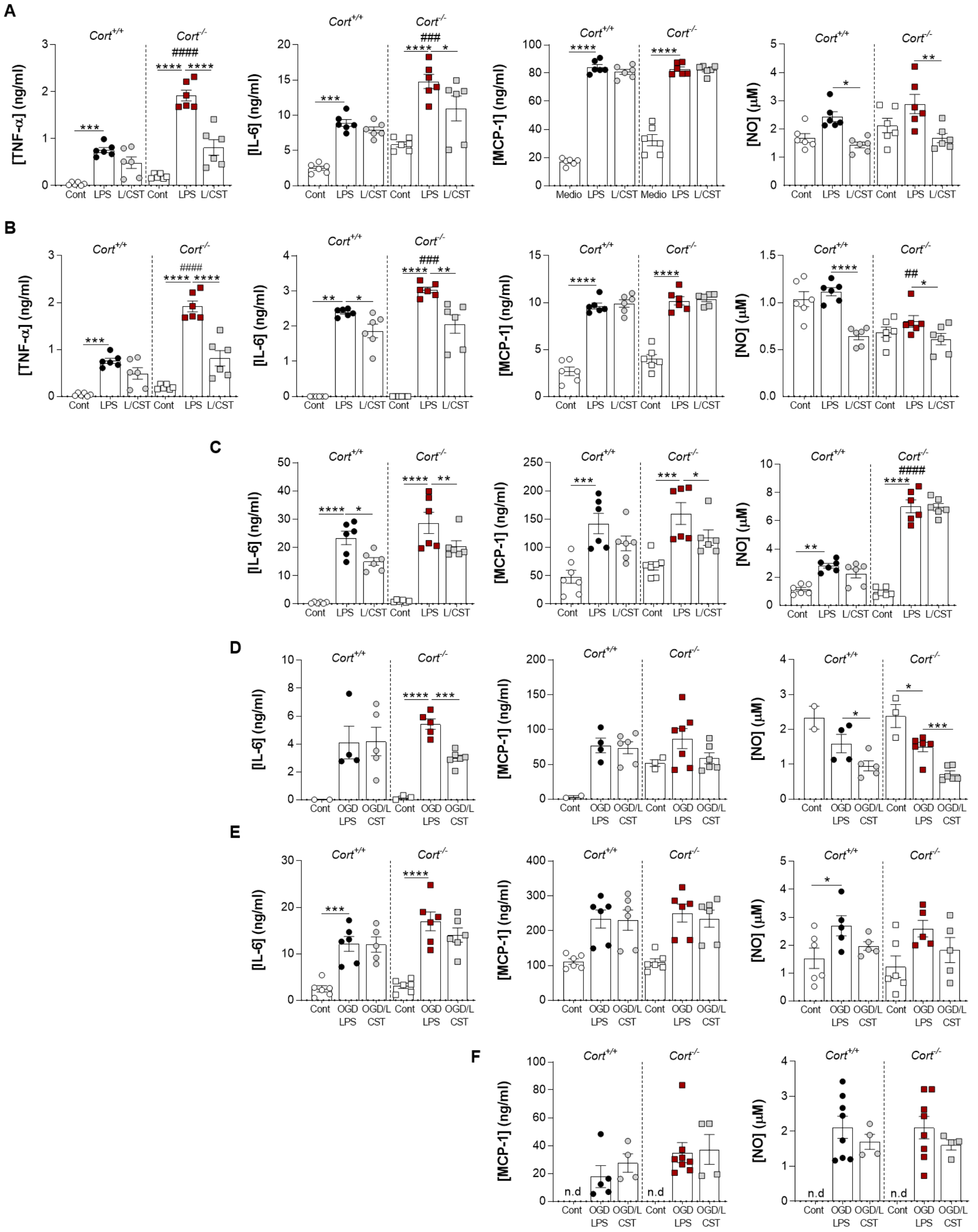


**Figure S8. Cortistatin is a key anti-inflammatory mediator in glial cells.** Primary cultures of astrocytes (**A, D**), mixed neuronal-glial cells (**B, E**), and microglia (**C, F**) were exposed to LPS (100 ng/ml) ± CST (100 nM) (**A–C**) or oxygen-glucose deprivation (2-4h) and reoxygenation (20-22h) followed by± CST and LPS (OGD/R + LPS ± CST) (**D–F**). Inflammatory markers were evaluated in the supernatants, with the results organized by columns: TNF-α (ng/mL, first column), IL-6 (ng/mL, second column), MCP-1 (ng/mL, third column), and nitric oxide (NO, µM, fourth column). **vs.* *Cort*$^{+/+}$ or *Cort*$^{-/-}$ LPS/OGD+LPS-treated cells; $^{\#}$*vs.* *Cort*$^{+/+}$ LPS/OGD+LPS-treated cells. $^{*/\#}p\leq0.05$. $^{**/\#\#}p\leq0.01$, $^{***/\#\#\#}p\leq0.001$, $^{****/\#\#\#\#}p\leq0.0001$. CST, cortistatin; OGD/L, oxygen-glucose deprivation + reperfusion +LPS.

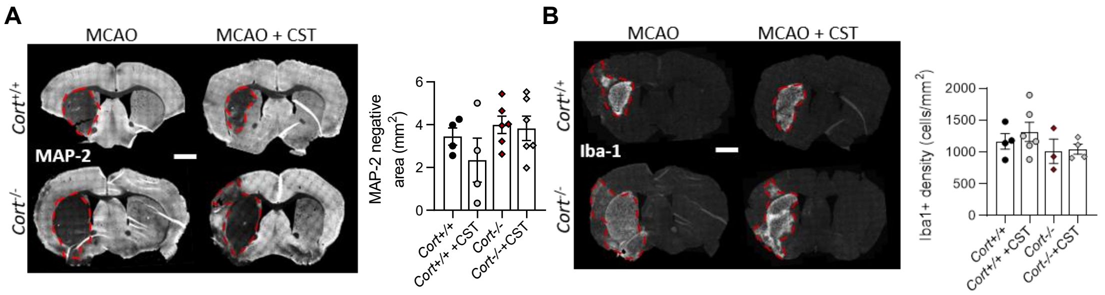


**Figure S9. Cortistatin modulates subacute stroke outcome**. Brain coronal sections were obtained from 12-weeks-old *Cort*$^{+/+}$ and *Cort*$^{-/-}$ mice treated daily with either saline (MCAO) or cortistatin (144 µg/kg) (MCAO+CST), from 24 h post-reperfusion, and sacrificed at 7 days. **(A)** Left, representative coronal sections stained with MAP-2. Ischemic lesions (loss of immunoreactivity/dark area) are delineated by dashed red lines. Scale bar: 1,500µm. Right, the absence of MAP-2 staining was quantified as a measure of neuronal damage. N=4-6 mice/group. **(B)** Left, representative sections stained with Iba1 to identify microglia/infiltrating macrophages. Right, Iba1+ density (cells/mm$^2$) was measured in the ischemic core. N=3-6 mice/group. Data represent the mean ± SEM. Statistical analyses were performed using unpaired t-tests or one-way ANOVA for comparisons among multiple groups. CST: cortistatin, Iba1: ionized calcium-binding adaptor molecule 1, MAP-2: microtubule-associated protein 2, MCAO: middle cerebral artery occlusion.

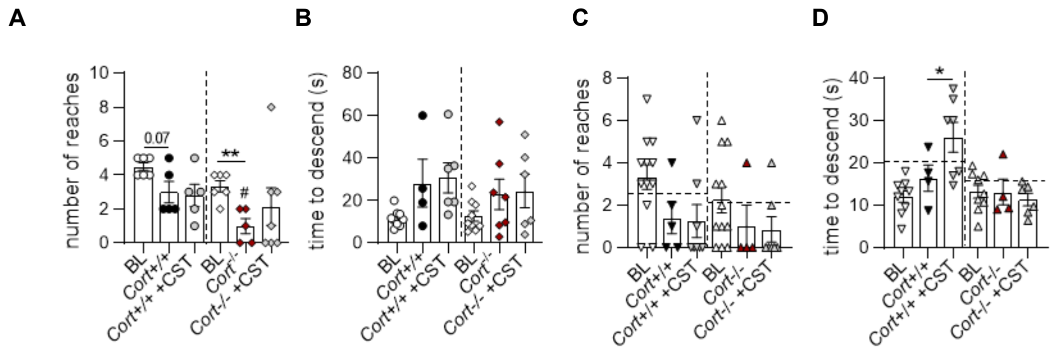


**Figure S10. Timely regulation of post-stroke outcomes by cortistatin.** 3-month-old **(A,B)** and 6-month-old (**C,D**) *Cort*$^{+/+}$ and *Cort*$^{-/-}$ mice were subjected to MCAO and received either saline or cortistatin (144μg/kg). Behavioural performances were assessed in 3-month-old animals at day 5 after stroke (treated daily with cortistatin) and 48 h post-surgery in 6-month-old animals (treated once with cortistatin 24 h post-reperfusion) using wire-hanging and pole tests, both at baseline (BL, pre-MCAO) and post-MCAO. In the wire-hanging test **(A,C)**, the number of reaches towards one of the posts was evaluated. 3-month-old animals (**A**): N=6 (*Cort*$^{+/+}$ and *Cort*$^{-/-}$ BL mice; MCAO+CST *Cort*$^{-/-}$ mice), N=5 (MCAO *Cort*$^{+/+}$ mice, MCAO+CST *Cort*$^{+/+}$ mice, MCAO *Cort*$^{-/-}$ mice); 6-month-old mice (**C**): N=12 (*Cort*$^{+/+}$ and *Cort*$^{-/-}$ BL mice), N=5 (MCAO *Cort*$^{+/+}$ mice), N=4 (MCAO *Cort*$^{-/-}$ mice), N=7 (MCAO+CST *Cort*$^{+/+}$ mice, MCAO+CST *Cort*$^{-/-}$ mice). In the pole test **(B,D)** the time to descend (s) was measured. 3-month-old animals **(B)**: N=9 (*Cort*$^{+/+}$ and *Cort*$^{-/-}$ BL mice), N=4 (MCAO *Cort*$^{+/+}$ mice), N=7 (MCAO *Cort*$^{-/-}$ mice), N=6 (MCAO+CST *Cort*$^{+/+}$ mice, MCAO+CST *Cort*$^{-/-}$ mice); 6-month-old mice **(D)**: N=9 (*Cort*$^{+/+}$ and *Cort*$^{-/-}$ BL mice), N=4 (MCAO *Cort*$^{+/+}$ mice, MCAO *Cort*$^{-/-}$ mice), N=7 (MCAO+CST *Cort*$^{+/+}$ mice), N=6 (MCAO+CST *Cort*$^{-/-}$ mice). Horizontal dashed lines represent values in 3-month-old *Cort*$^{+/+}$ (black) and *Cort*$^{-/-}$ (red) mice subjected to MCAO and treated with saline. Data are the mean ± SEM with dots representing individual mice. Statistical analyses were performed using unpaired t-tests or one-way ANOVA for comparisons among multiple groups. **vs. Cort*$^{+/+}$ or *Cort*$^{-/-}$ MCAO saline-treated mice*;* **p* ≤ 0.05. CST: cortistatin

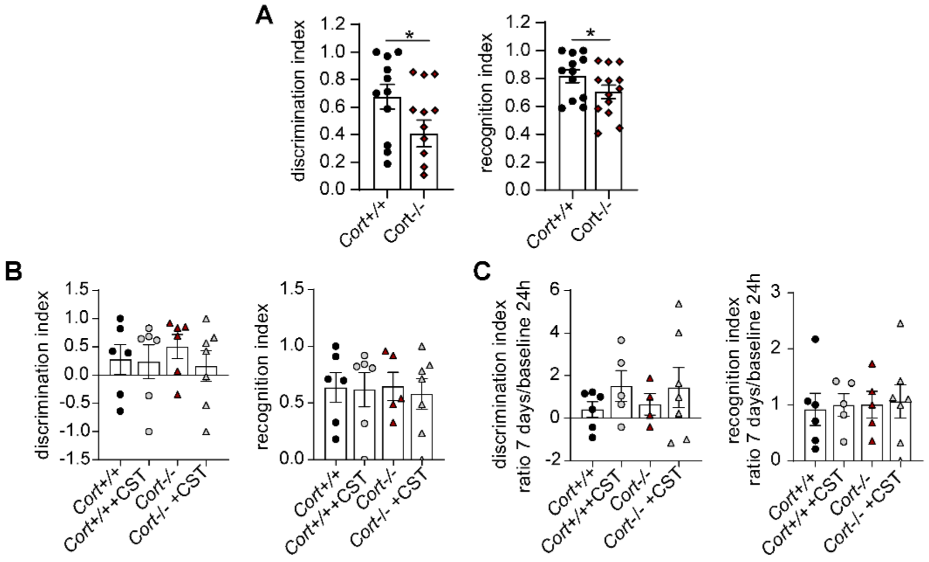


**Figure S11. Cortistatin deficiency is associated with cognitive impairment.** *Cort*$^{+/+}$ and *Cort*$^{-/-}$ mice underwent MCAO and received daily treatments of saline or cortistatin (144 µg/kg) from 24 h post-reperfusion until day 7. Memory was assessed using the novel object recognition test. Performance was evaluated at baseline (pre-stroke) (A) and 7 days post-stroke (B). During the evaluations, mice were exposed to two identical objects, and 24 h later, one was replaced by a novel object. Exploration time (t, snout ≤ 2 cm from the object) was recorded over a 10-min trial using SMART automated tracking. Memory performance was quantified using the discrimination index ([t-novel - t-familiar] / [t-novel + t-familiar]; left graphs), the recognition index (t-novel / [t-novel + t-familiar]; right graphs, or the ratio of these indexes between 7 days and baseline (C). N = 6–7 mice per group. Data are mean ± SEM, with dots representing individual mice. Statistical analyses were performed using unpaired t-tests or one-way ANOVA. *vs. *Cort-/-* MCAO mice; *$p \leq 0.05$.

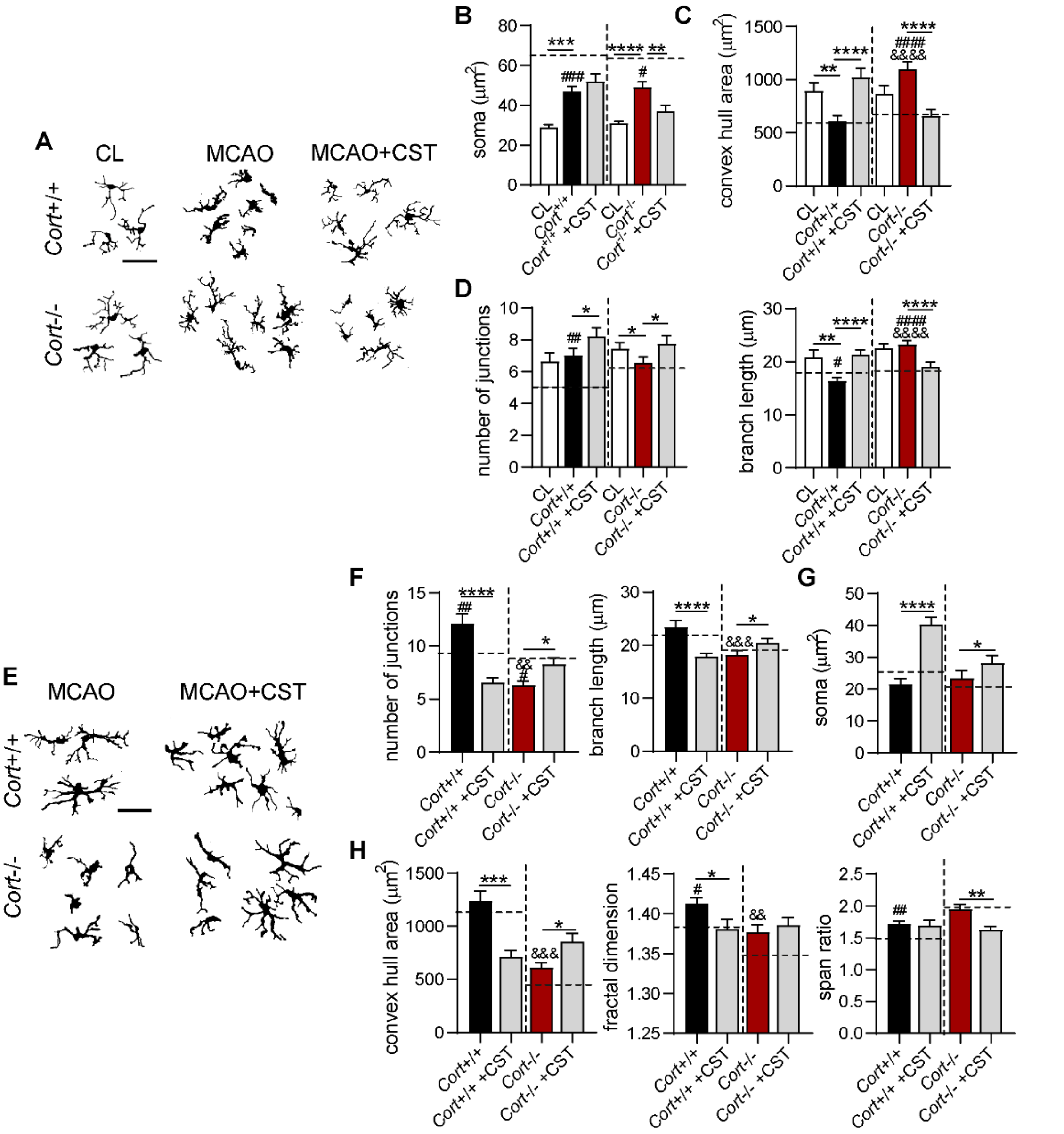


**Figure S12. Lack of cortistatin exacerbates glial dysregulation in 6-month-old mice.** 6-month-old $Cort^{+/+}$ and $Cort^{-/-}$ mice were subjected to MCAO and treated with either saline or cortistatin (144µg/kg) at 24 h post-reperfusion and sacrificed at 48 h. **(A)** Binary images of Iba1+ cells from CL and peri-lesion. Scale bar: 30µm. **(B-D)** Quantification of $Iba1^{+}$ microglial morphology: soma size, manually measured **(B)**, convex hull area, evaluated by fractal analysis **(C)**, and junction number and branch length, determined by skeleton analysis **(D)**. Cell counts per group: n=30-35 cells, N=3 mice/group. **(E).** Representative binary images of glial scar astrocytes. Scale bar: 30µm. **(F-H).** Astrocyte morphological features analysed via skeleton and fractal analysis: junctions and branching, **(F)**, soma size **(G)**, and convex hull area, fractal dimension and span ratio **(H)**. Cell counts per group: n=30-35 cells, N=3 mice/group. Data were compared to MCAO saline-treated 3-month-old $Cort^{+/+}$ and $Cort^{-/-}$ mice (dashed line). Data are expressed as mean ± SEM. Statistical analyses were performed using unpaired t-tests or one-way ANOVA for comparisons among multiple groups. $^{*}$*vs.* $Cort^{+/+}$ or $Cort^{-/-}$ MCAO saline-treated mice*;* $^{\#}$*vs.* 3-month-old $Cort^{+/+}$ or $Cort^{-/-}$ MCAO saline-treated mice*;* $^{\&}$*vs.* 6-month-old $Cort^{+/+}$ MCAO saline-treated mice $^{*/\#/\&}p\leq0.05$. $^{*/\#/\&}p\leq0.05$, $^{**/\#\#/\&\&}p\leq0.01$, $^{***/\#\#\#/\&\&\&}p\leq0.001$, $^{****/\#\#\#\#/\&\&\&\&}p\leq0.0001$. CST, cortistatin; MCAO, middle cerebral artery occlusion.

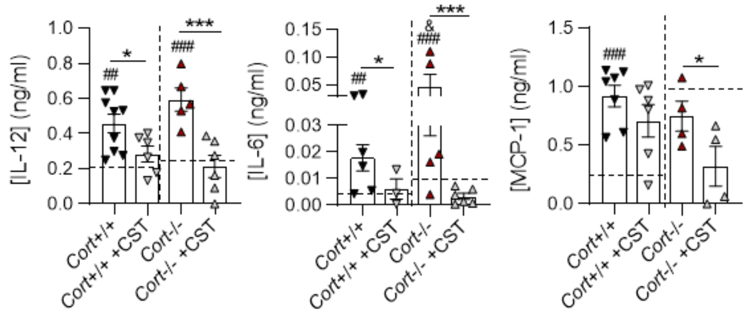


**Figure S13. Cortistatin alleviates peripheral immune activation in 6-month-old mice suffering stroke.** Serum levels of IL-12, IL-6, and MCP-1 (ng/ml) were measured using an ELISA assay in 6-month-old MCAO mice 48 h after ischemic stroke. IL-12: N=9 (MCAO *Cort*$^{+/+}$ mice), 6 (MCAO+CST *Cort*$^{+/+}$ mice), 5 (MCAO *Cort*$^{-/-}$ mice), 6 (MCAO+CST *Cort*$^{-/-}$ mice); IL-6: N=6 (MCAO *Cort*$^{+/+}$ mice), 3 (MCAO+CST *Cort*$^{+/+}$ mice), 5 (MCAO *Cort*$^{-/-}$ mice), 6 (MCAO+CST *Cort*$^{-/-}$ mice); MCP-1: N=7 (MCAO *Cort*$^{+/+}$ mice), 6 (MCAO+CST *Cort*$^{+/+}$ mice), 4 (MCAO *Cort*$^{-/-}$ mice), 4 (MCAO+CST *Cort*$^{-/-}$ mice). Dashed lines represent values in 3-month-old *Cort*$^{+/+}$ and *Cort*$^{-/-}$ mice subjected to MCAO and treated with saline. Data are the mean ± SEM with dots representing individual mice. Statistical analyses were performed using unpaired t-tests or one-way ANOVA for comparisons among multiple groups. **vs*. 6-month-old *Cort*$^{+/+}$ or *Cort*$^{-/-}$ MCAO saline-treated mice; $^{\#}$*vs.* 3-month-old *Cort*$^{+/+}$ or *Cort*$^{-/-}$ MCAO saline-treated mice; $^{\&}$vs. 6-month-old *Cort*$^{+/+}$ MCAO saline-treated mice. $^{*/\#/\&}p \leq 0.05$. $^{*/\#/\&}p \leq 0.05$, $^{**/\#\#/\&\&}p \leq 0.01$, $^{***/\#\#\#/\&\&\&}p \leq 0.001$, $^{****/\#\#\#\#/\&\&\&\&}p \leq 0.0001$.CST: cortistatin, IL-6: interleukin 6, IL-12: interleukin 12, MCAO: middle cerebral artery occlusion, MCP-1: Monocyte Chemoattractant Protein- 1.

### C. Extended References


Andrés-León, E., and A.M. Rojas. 2019. miARma-Seq, a comprehensive pipeline for the simultaneous study and integration of miRNA and mRNA expression data. *Methods*. 152:31–40. doi:10.1016/j.ymeth.2018.09.002.

Androvic, P., D. Kirdajova, J. Tureckova, D. Zucha, E. Rohlova, P. Abaffy, J. Kriska, M. Valny, M. Anderova, M. Kubista, and L. Valihrach. 2020. Decoding the Transcriptional Response to Ischemic Stroke in Young and Aged Mouse Brain. *Cell Rep*. 31:107777. doi:10.1016/j.celrep.2020.107777.

Brionne, A., A. Juanchich, and C. Hennequet-Antier. 2019. ViSEAGO: a Bioconductor package for clustering biological functions using Gene Ontology and semantic similarity. *BioData Min*. 12:16. doi:10.1186/s13040-019-0204-1.

Castanza, A.S., J.M. Recla, D. Eby, H. Thorvaldsdóttir, C.J. Bult, and J.P. Mesirov. 2023. Extending support for mouse data in the Molecular Signatures Database (MSigDB). *Nat Methods*. 20:1619–1620. doi:10.1038/s41592-023-02014-7.

Clough, E., T. Barrett, S.E. Wilhite, P. Ledoux, C. Evangelista, I.F. Kim, M. Tomashevsky, K.A. Marshall, K.H. Phillippy, P.M. Sherman, H. Lee, N. Zhang, N. Serova, L. Wagner, V. Zalunin, A. Kochergin, and A. Soboleva. 2024. NCBI GEO: archive for gene expression and epigenomics data sets: 23-year update. *Nucleic Acids Res*. 52:D138–D144. doi:10.1093/nar/gkad965.

Huber, W., V.J. Carey, R. Gentleman, S. Anders, M. Carlson, B.S. Carvalho, H.C. Bravo, S. Davis, L. Gatto, T. Girke, R. Gottardo, F. Hahne, K.D. Hansen, R.A. Irizarry, M. Lawrence, M.I. Love, J. MacDonald, V. Obenchain, A.K. Oleś, H. Pagès, A. Reyes, P. Shannon, G.K. Smyth, D. Tenenbaum, L. Waldron, and M. Morgan. 2015. Orchestrating high-throughput genomic analysis with Bioconductor. *Nat Methods*. 12:115–121. doi:10.1038/nmeth.3252.

Mancarci, B.O., L. Toker, S.J. Tripathy, B. Li, B. Rocco, E. Sibille, and P. Pavlidis. 2017. Cross-Laboratory Analysis of Brain Cell Type Transcriptomes with Applications to Interpretation of Bulk Tissue Data. *eNeuro*. 4:ENEURO.0212-17.2017. doi:10.1523/ENEURO.0212-17.2017.

Naba, A., K.R. Clauser, S. Hoersch, H. Liu, S.A. Carr, and R.O. Hynes. 2012. The Matrisome: In Silico Definition and In Vivo Characterization by Proteomics of Normal and Tumor Extracellular Matrices. *Mol & Cell Proteomics*. 11. doi:10.1074/mcp.M111.014647.

Nikolayeva, O., and M.D. Robinson. 2014. edgeR for Differential RNA-seq and ChIP-seq Analysis: An Application to Stem Cell Biology. *In* Stem Cell Transcriptional Networks: Methods and Protocols. B.L. Kidder, editor. Springer, New York, NY. 45–79.

Reeb, P.D., S.J. Bramardi, and J.P. Steibel. 2015. Assessing Dissimilarity Measures for Sample-Based Hierarchical Clustering of RNA Sequencing Data Using Plasmode Datasets. *PLoS One*. 10:e0132310. doi:10.1371/journal.pone.0132310.

Ritchie, M.E., B. Phipson, D. Wu, Y. Hu, C.W. Law, W. Shi, and G.K. Smyth. 2015. limma powers differential expression analyses for RNA-sequencing and microarray studies. *Nucleic Acids Res*. 43:e47. doi:10.1093/nar/gkv007.

Schep, A.N., and S.K. Kummerfeld. 2017. iheatmapr: Interactive complex heatmaps in R. *J Open Source Softw*. 2:359. doi:10.21105/joss.00359.

Subramanian, A., P. Tamayo, V.K. Mootha, S. Mukherjee, B.L. Ebert, M.A. Gillette, A. Paulovich, S.L. Pomeroy, T.R. Golub, E.S. Lander, and J.P. Mesirov. 2005. Gene set enrichment analysis: A knowledge-based approach for interpreting genome-wide expression profiles. *Proc Natl Acad Sci U S A*. 102:15545–15550. doi:10.1073/pnas.0506580102.

Wickham, H. 2011. ggplot2. *WIREs Comput Stat*. 3:180–185. doi:10.1002/wics.147.

Wickham, H. 2016. ggplot2: Elegant Graphics for Data Analysis. Springer-Verlag New York.

Yu, G., L.-G. Wang, Y. Han, and Q.-Y. He. 2012. clusterProfiler: an R package for comparing biological themes among gene clusters. *OMICS*. 16:284–287. doi:10.1089/omi.2011.0118.

Zheng, K., L. Lin, W. Jiang, L. Chen, X. Zhang, Q. Zhang, Y. Ren, and J. Hao. 2022. Single-cell RNA-seq reveals the transcriptional landscape in ischemic stroke. *J Cereb Blood Flow Metab*. 42:56–73. doi:10.1177/0271678X211026770.
